# Selective binders of a cotranslational folding intermediate

**DOI:** 10.64898/2026.10.30.763055

**Authors:** Ohm Prakash, Tomas B. Voisin, Alžběta Roeselová, Cecilie M. Jørgensen, James S. Desmet, Andrea Nans, Rick Davies, Louise J. Walport, David Balchin

**Author notes:** Correspondence to David Balchin, or Louise Walport.

## Abstract

Drug discovery efforts typically target mature, folded proteins, primarily to inhibit their function or induce their degradation. An alternative strategy would be to target incompletely folded states that are transiently populated during protein biogenesis, but whether these states contain druggable pockets was unclear. Here, we used an mRNA display approach to discover cyclic peptides that recognize a nascent polypeptide at an intermediate point during its synthesis on the ribosome. We identified peptides that bind with low-nanomolar affinity to a model nascent protein, β-galactosidase, but do not detectably bind either the fully-folded protein or ribosomes synthesising different sequences. Hydrogen/deuterium exchange-mass spectrometry and cryogenic electron microscopy revealed the molecular basis for selective binding. We found that the nascent chain folds into an α-helix against the ribosome surface during ongoing translation, creating a cryptic pocket together with ribosomal proteins uL23 and uL29 that is exploited by the hit peptides. Our results establish proof-of-concept for selective targeting of protein biogenesis intermediates, and provide a platform for future drug development.

## Introduction

Most drugs are small molecules that target mature disease-causing proteins to modulate their activity. For difficult targets, non-traditional modalities such as cyclic peptides and antibodies are also being pursued, due to their ability to bind larger, less well-defined protein surfaces^1^. Alternatively, more recent approaches have involved reducing protein levels, either through targeting specific mRNA^2^, or by inducing degradation by means of a PROTAC (proteolysis-targeting chimera)^3^ or another bifunctional molecule^4^. However, many disease-relevant protein targets remain out of reach of current drug modalities and screening strategies. This is especially true for proteins that lack obvious binding pockets, such as those involved in protein-protein interactions, that are intrinsically disordered, or adopt in vivo conformations, such as aggregates, that mask viable binding sites^5^.

The process of protein synthesis may offer new possibilities for drugging classically “undruggable” proteins. Ribosome-targeting antibiotics, such as chloramphenicol and erythromycin, have been used for over a century to kill bacteria by halting all cellular translation in a non-specific manner^6^. More recently, several small molecules have been reported, that bind deep in the ribosomal exit tunnel and make additional contacts with several amino acids in the nascent protein to provide some protein-level selectivity^7–9^. We hypothesized that greater selectivity might be achieved by targeting nascent proteins as they emerge from the ribosome. Proteins begin to fold and assemble during their synthesis on the ribosome, forming stable cotranslational folding intermediates that often differ from those populated post-translation^10–15^ (Fig. 1a). In addition, these partially-synthesized intermediates expose incomplete domain and subunit interfaces, potentially providing new binding sites that are accessible only during translation^16,17^. Binding to cotranslational folding intermediates is expected to down-regulate the target protein by triggering stringent cellular quality control pathways that monitor translating ribosomes. Indeed, interfering with ribosome processivity or nascent protein folding has been shown to lead to selective degradation of the defective nascent protein and its encoding mRNA^18–23^.

**Fig. 1.**
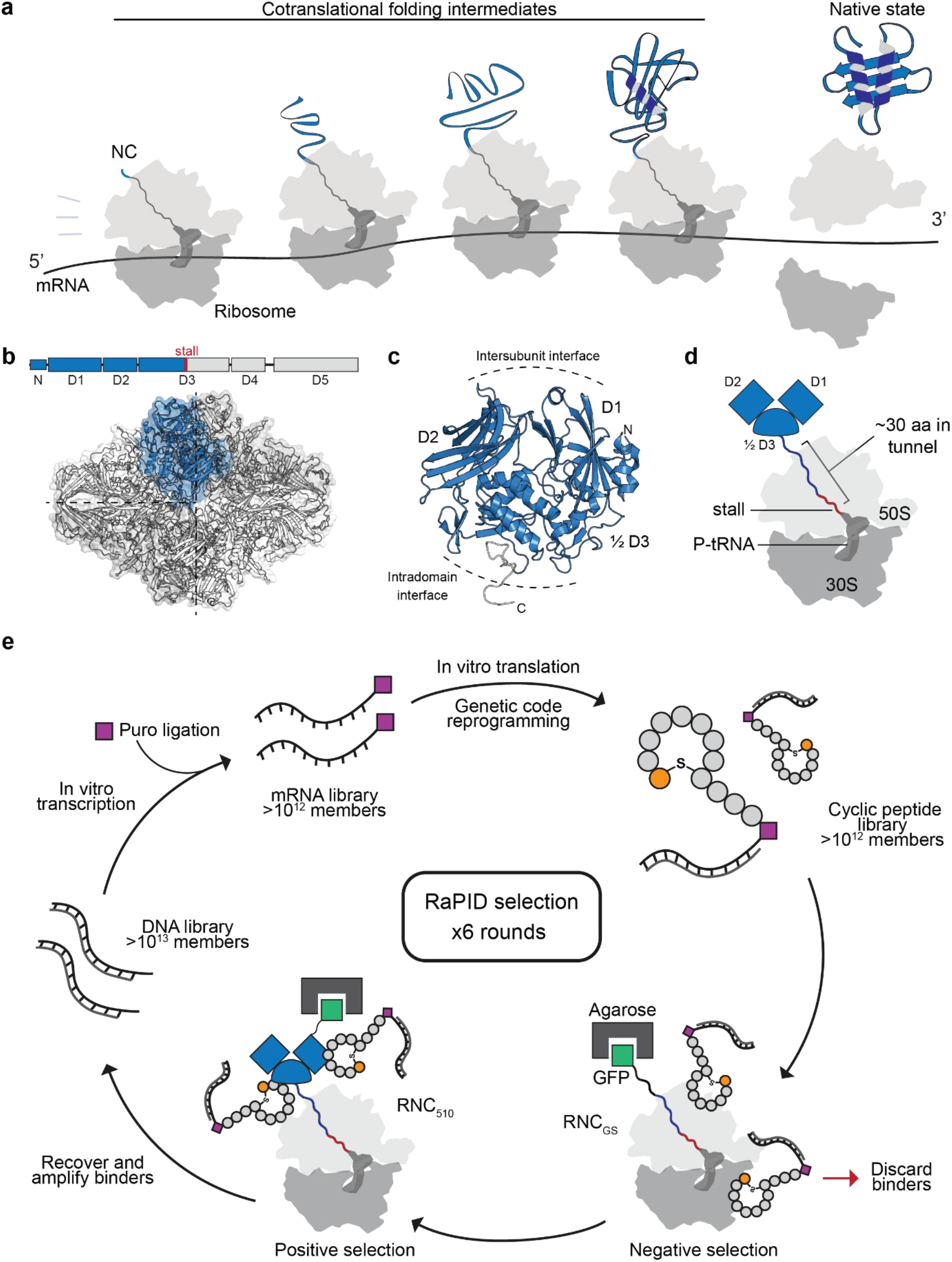
mRNA display applied to cotranslational folding intermediates. a, Proteins begin to fold during their synthesis on the ribosome and temporarily populate partially-folded intermediate states. **b**, A partially-synthesised fragment of β-gal as the target for binder discovery. Top, domain organization of *E. coli* β-gal showing the position at which translation is stalled, after synthesis of residues 1-510. Bottom, cartoon representation of native tetrameric β-gal, with residues 1-510 in one monomer colored blue (PDB: 6CVM^24^). Subunit interfaces are indicated with dashed lines. **c**, Cartoon representation of the native structure of residues 1-510 of β-gal. **d**, Schematic of RNC510. **e**, RaPID (Random non-standard Peptides Integrated Discovery) workflow for identifying cyclic peptide binders of RNC510. A large semi-randomized DNA library is transcribed and ligated to puromycin (puro), before in vitro translation and genetic code reprogramming to introduce N-chloroacetylated tryptophan (ClAcW) in place of the initiator methionine (orange). ClAcW spontaneously reacts with a proximal cysteine residue, resulting in a cyclic peptide covalently linked (via puro) to its encoding mRNA. The peptide library is first added to immobilized RNCGS to remove non-specific binders of the ribosome or muGFP, before being applied to immobilized RNC510. Binders are recovered, amplified by PCR and enriched over multiple rounds of selection.

Here, we establish proof-of-concept for selective targeting of cotranslational folding intermediates. We demonstrate that ribosome-tethered folding intermediates contain druggable pockets, and detail a generalizable approach to discover cyclic peptides that bind these pockets with high affinity. For a model bacterial protein, β-galactosidase, we identify cotranslational binders that exploit a specific neo-interface formed by the partially-folded nascent polypeptide and the ribosome surface. Overall, our study introduces a new approach to expand druggable space for challenging targets.

## Results

### RaPID mRNA display applied to ribosome-tethered folding intermediates

We first sought to develop a platform to discover selective binders of cotranslational folding intermediates. To establish proof-of-concept, we chose *E. coli* β-galactosidase (β-gal) as a well-characterized model system. β-gal is a large tetrameric enzyme with each subunit consisting of five structured domains and a disordered N-terminal region^25^ (Fig. 1b). We previously showed that β-gal folds and assembles cotranslationally, and that this is critical for its efficient maturation^11^. As the target for cotranslational binder discovery, we prepared a stalled ribosome:nascent chain complex (RNC) containing the first 510 residues of β-gal trapped on the ribosome (RNC_510_, Fig. 1b-d). We generated RNC_510_ by expressing a stalling construct in *E. coli*, consisting of an N-terminal muGFP (monomeric ultrastable green fluorescent protein), residues 1-510 of β-gal, and a 10-residue stalling sequence at the C-terminus^11,26,27^. Synthesis of the penultimate residue of the stalling sequence by endogenous *E. coli* ribosomes distorts the peptidyl transfer centre and prevents formation of the subsequent peptide bond^28^. The resulting RNCs consist of intact 70S ribosomes stably tethered to the NC via peptidyl-tRNA (Fig. 1d). In RNC_510_, the N-terminal 2½ domains of β-gal are exposed on the ribosome and the NC is partially folded and unassembled, thus exposing potential binding pockets that are inaccessible in the mature tetramer (Fig. 1c,d).

To discover binders of RNC_510_, we screened a pooled peptide macrocycle library using RaPID (Random non-standard Peptides Integrated Discovery, Fig. 1e)^29–32^. We used a large DNA library (>10^13^ members) to produce semi-randomized peptide sequences of the form ClAcW(X)_6-12_CGSGSGS, where the initiator methionine was reprogrammed to an N-chloroacetylated tryptophan (ClAcW) that spontaneously cyclises with the C-terminal cysteine, and X can be any amino acid. The DNA library was transcribed to RNA, conjugated to puromycin, reverse transcribed and translated to cyclic peptides by in vitro translation (IVT).

The 3’-puromycin ensures that each translated peptide is covalently linked to the mRNA template that encoded it. We immobilized RNC_510_ to agarose beads via the N-terminal GFP tag and performed successive rounds of affinity panning, identifying enriched peptides by sequencing the cDNA of each ligated mRNA. To produce a screening library depleted of sequences that bind non-specifically to the ribosome or GFP, we counter-screened the peptide library (from rounds 2-6) using RNC_GS_, in which the β-gal sequence was replaced with a 40-residue GlySer-rich linker.

### Identification of cyclic peptides that bind selectively to nascent β-gal

After 6 rounds of RaPID, we selected the top 1000 most enriched sequences and clustered these based on sequence homology^33,34^ (Extended Data Figures 1 and 2). Three prominent clusters emerged, and we focused on the most enriched peptide from each, hereafter named Peptide A, B and C (Fig. 2a,b). Peptides A (*N*-AcWTEFIRTC(S-)LWNHAGS-*C*) and B (*N*-AcWTRYIRTC(S-)HWVAGS-*C*) were similar in sequence, differing at only 4 positions. Both peptides featured a cysteine at position 8 and were synthesized as macrocycles closed at this position, followed by a C-terminal tail of 6 or 7 residues in which the second cysteine encoded in the enriched sequence was replaced by an alanine. Peptide C (*N*-AcWNWVLIRYGLKKC(S-)GS-*C*) was synthesized as a larger macrocycle closed at position 13.

**Fig. 2.**
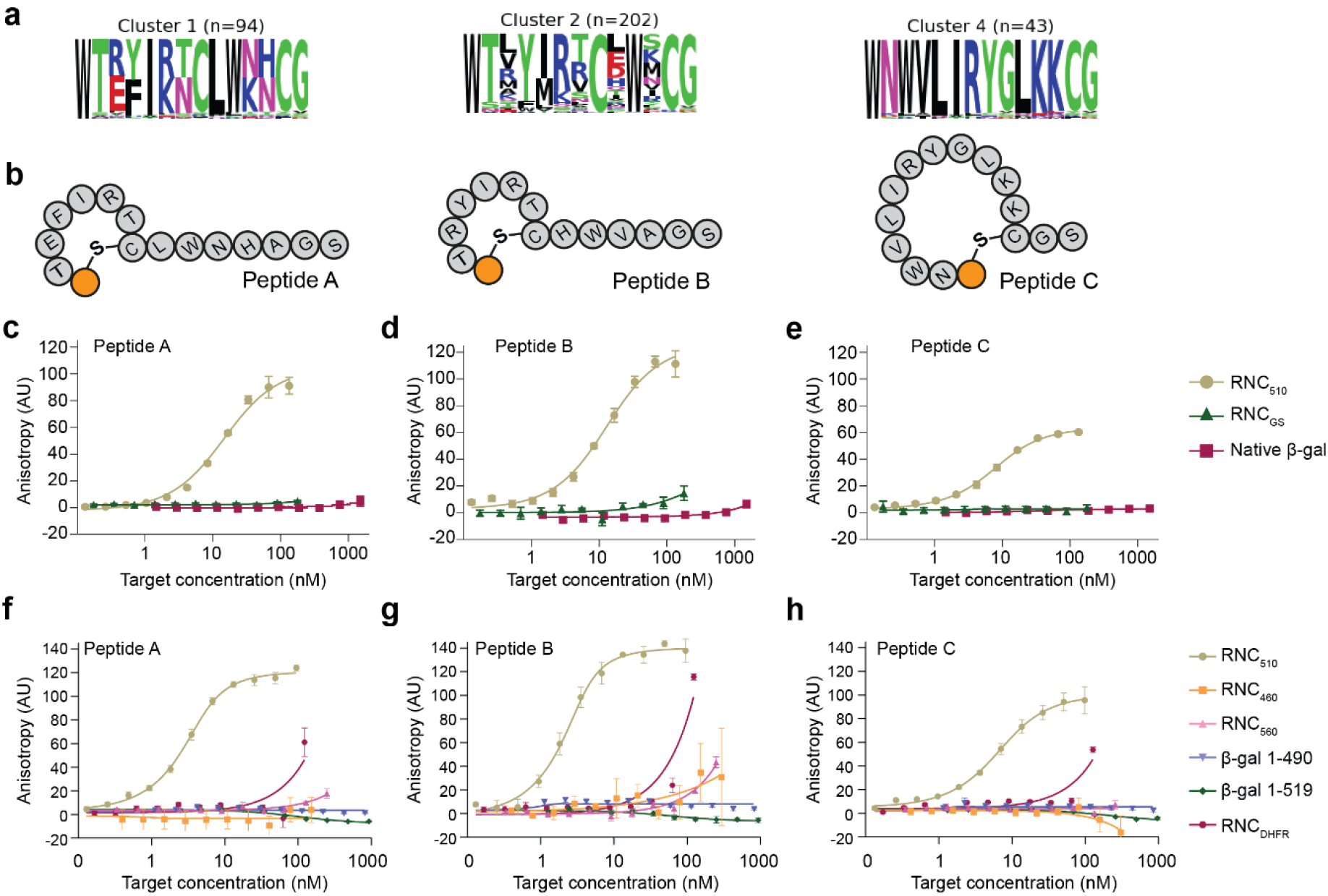
Tight and specific binders of nascent β-gal on the ribosome. a, Logo plots for prominent sequence clusters. **b**, Schematics of representative peptides chosen from each cluster for synthesis. **c-h,** Quantification of peptide binding. Carboxyfluorescein-labelled peptide was incubated with increasing concentrations of different RNCs or off-ribosome β-gal variants, and binding was measured by the increase in fluorescence anisotropy. Data are presented as mean ± SD, n = 3 independent experiments.

We prepared variants of each peptide with a C-terminal carboxyfluorescein, and quantified binding to RNCs using fluorescence anisotropy (Fig. 2c-e). Each peptide bound tightly to RNC_510_, with dissociation constants of 12 ± 2 nM (Peptide A), 10 ± 1 nM (Peptide B) and 5.3 ± 0.9 nM (Peptide C). In contrast, we observed negligible or very weak binding to RNC_GS_, or RNC_DHFR_ exposing residues 1-126 of *E. coli* dihydrofolate reductase^10^, demonstrating that the peptides are selective for ribosomes translating β-gal (Fig. 2f-h). Moreover, the peptides were selective for the stage of β-gal synthesis, as they did not bind RNCs exposing shorter (RNC_460_) or longer (RNC_560_) NCs (Fig. 2f-h). The context of the ribosome was also required for binding, as we did not detect binding to full-length tetrameric β-gal, nor to soluble off-ribosome fragments of β-gal of similar length to the NC (1-519 and 1-490)^35^ (Fig. 2c-h).

In summary, we identified cyclic peptides that bind specifically and with high affinity to nascent β-gal on the ribosome. The peptides recognize features of the intact β-gal RNC, rather than the ribosome or NC in isolation, and they distinguish the NC from mature tetrameric β-gal.

### Cyclic peptides bind an intermediate formed during ongoing translation

Since the peptides were selected for their ability to bind a stalled RNC, we next sought to establish whether they also bind during ongoing β-gal synthesis. We programmed a fully-reconstituted IVT system using mRNA encoding full-length β-gal or DHFR as a negative control, and assayed peptide binding by fluorescence anisotropy as above (Fig. 3a). For all three peptides, we observed a strong increase in anisotropy ∼10 min after supplying the IVT with β-gal mRNA, indicative of peptide binding (Fig. 3b-d). This timing is consistent with peptide binding to an incompletely-synthesised folding intermediate, since synthesis of full-length β-gal requires ∼15 min under identical conditions^11^. To test whether the peptides selectively bound nascent β-gal in the complex transcription/translation reaction mixture, we synthesized a variant of peptide C containing a photo-leucine residue for photo-crosslinking. IVTs programmed with mRNA encoding β-gal, but not DHFR, showed a crosslinked product consistent in size with the nascent protein (Extended Data Fig. 3a,b). To determine whether specific sequence elements in the NC are required for binding, we expressed truncated variants of β-gal. Constructs encoding the first 3 or 4 domains of β-gal (1-667, 1-745) bound and crosslinked to the peptides, whereas constructs encoding fragments shorter than 2½ domains (1-332, 1-440, 1-490) did not (Fig. 3b-d and Extended Data Fig. 3a,b).

**Fig. 3.**
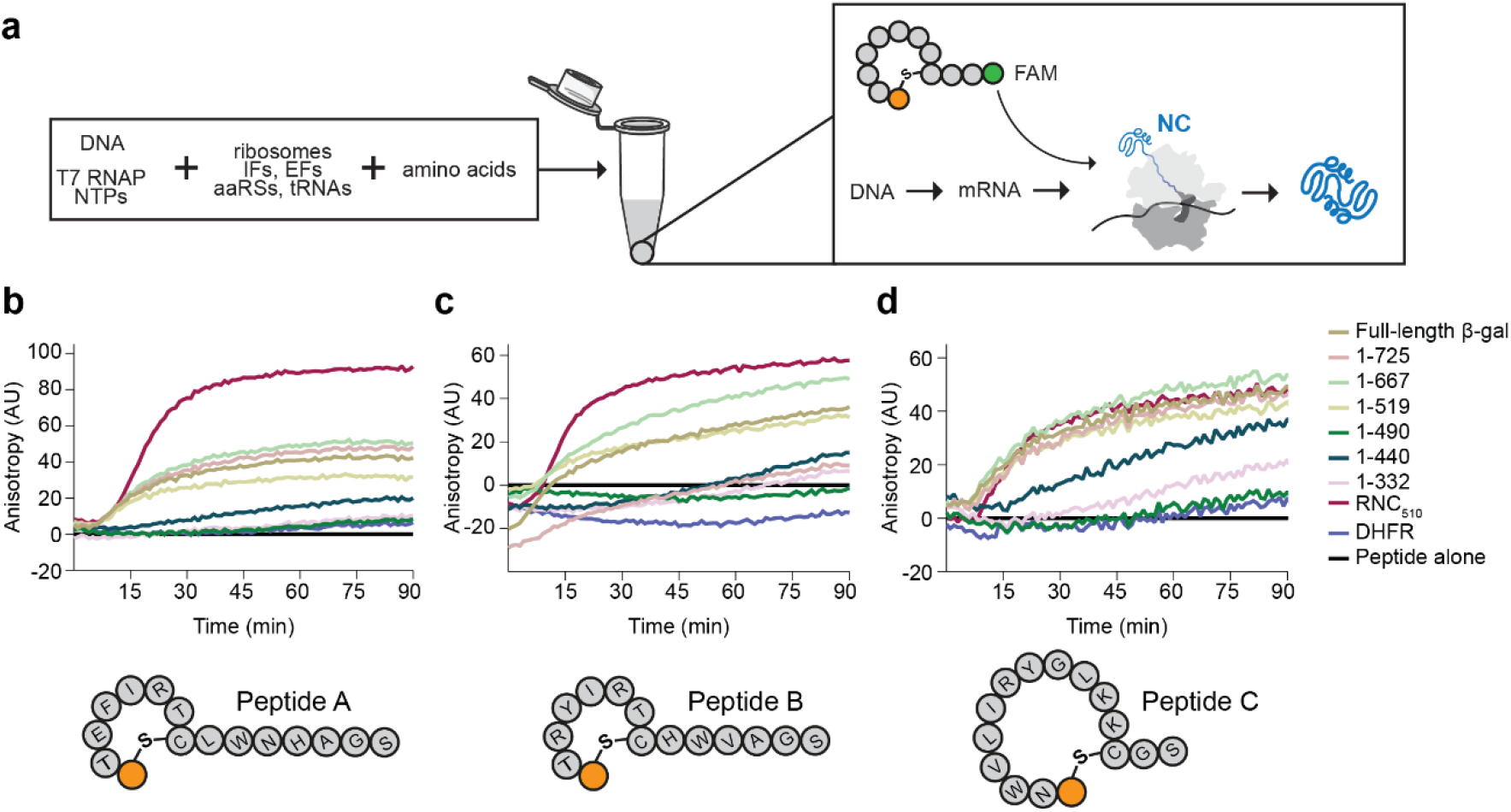
Cyclic peptides bind an intermediate formed during ongoing translation. a, Schematic of reconstituted in vitro transcription/translation system for measuring peptide binding during real-time translation. **b-d**, Binding of FAM-labelled peptides during synthesis of full-length β-gal, β-gal truncations, full-length DHFR or β-gal 1-510 with a C-terminal ribosome stalling sequence (RNC510) was measured by fluorescence anisotropy.

Together, these data demonstrate that the peptides bind an intermediate of β-gal formed during ongoing translation, and that they recognise sequence or structural elements near the C-terminus of the NC in RNC_510_.

### Cyclic peptides recognize a composite pocket formed by the nascent protein and ribosome surface

We next sought to identify the peptide binding site(s) on RNC_510_. Partially-synthesised NCs are conformationally dynamic and typically cannot be resolved by standard structural biology approaches. Therefore, we first assessed peptide binding using hydrogen/deuterium exchange coupled to mass spectrometry (HDX-MS). In HDX-MS, molecular interactions and the resulting allosteric conformational changes are detected as changes in the rate of amide hydrogen-to-deuterium exchange, quantified for all proteins in the system by MS^36^. We used a bespoke HDX-MS workflow, tailored to large and RNA-rich RNCs, to compare peptide-level deuteron incorporation in the presence and absence of each peptide^26^ (Fig 4a). To capture dynamics at different timescales, we exposed RNCs to deuterium for 10, 100 or 3000 s. We followed 53 β-gal peptides in each condition, covering 76% of the NC sequence (Data S2). The pattern of protection from deuterium exchange was similar for all 3 peptides, suggesting a common mode of NC engagement (Fig. 4b-d, Extended Data Fig. 4a-f and Data S2). The protected sites clustered near the C-terminus of the NC, with the most strongly protected region (residues 472-497) mapping to the part of nascent β-gal closest to the exit port of the ribosome (Fig. 4e). To determine whether the peptides also bind the ribosome itself, we measured deuteron incorporation in ribosomal proteins uL23 and uL29 that form part of the vestibule. Binding of all 3 peptides protected part of uL29 directly at the exit port, while peptides A and B additionally protected part of uL23 that abuts uL29 (Fig. 4f-l, Extended Data Fig. 5a-l and Data S1). Together, our HDX-MS analysis suggested that the peptides bind NC elements close to the ribosome exit port, and in addition directly contact the ribosome surface.

**Fig. 4.**
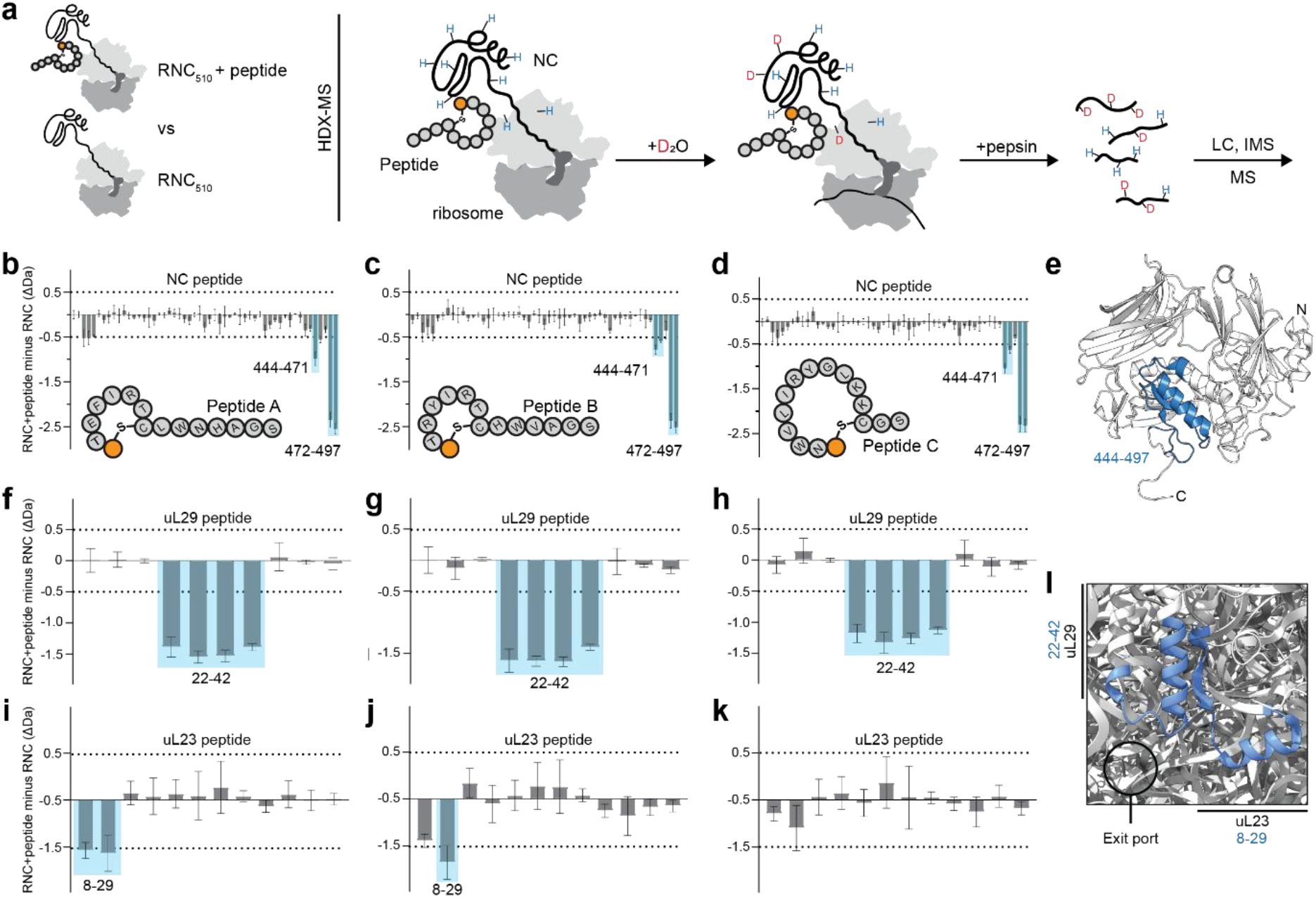
Peptides contact ribosomal proteins and the nascent chain near the exit port. a, Schematic of HDX-MS experiment. **b-d**, Protection of β-gal NC peptides induced by cyclic peptide binding. Difference in deuteron uptake after 10 s, between RNC510 alone and in complex with each cyclic peptide. Negative values indicate protection from exchange in the presence of the peptide. **e**, Native structure of β-gal 1-510, with residues protected by peptide binding colored blue. **f-k**, Protection of uL23 or uL29 peptides induced by cyclic peptide binding. Difference in deuteron uptake after 100 s, between RNC510 alone and in complex with each cyclic peptide. Negative values indicate protection from exchange in the presence of the peptide. **l**, Structure of uL23 and uL29 at the ribosome exit port, with residues protected by peptide binding colored blue (PDB: 7K00^37^). HDX-MS data are presented as mean ± SD, n = 3 independent labelling reactions. Uptake data for all peptides are shown in Data S2.

We next determined the structures of RNC_510_ bound to each peptide using single-particle cryogenic electron microscopy (cryoEM). The reconstructions reached a global resolution of 2.2 to 2.4 Å and showed no significant difference in the ribosome density compared to previous structures of stalled RNCs^11^ (Extended Data Fig. 6-8 and Extended Data Table 2). As in our previous cryoEM structure of apo RNC_510_, residues 466 to 478 of the NC form a stable helix on the surface of the ribosome between uL23 and uL29, while residues 1-478 are unresolved^11^ (Fig. 5a-d). In addition to the NC helix, we observed clear density for the bound peptides. Consistent with our HDX-MS analysis, all peptides bind near the ribosomal exit port and make contacts with the nascent chain as well as ribosomal proteins uL23 and uL29 (Fig. 5a-e). The peptides showed well-defined density corresponding to their cycles, although their C-tails were not fully resolved (Fig 5b-d). Peptides A and B, whose cyclic portions differ by only two residues, adopt almost identical binding poses. Peptide C binds the RNC in a different orientation, and part of the cycle folds into a short α-helix (Fig. 5d,e).

**Fig. 5.**
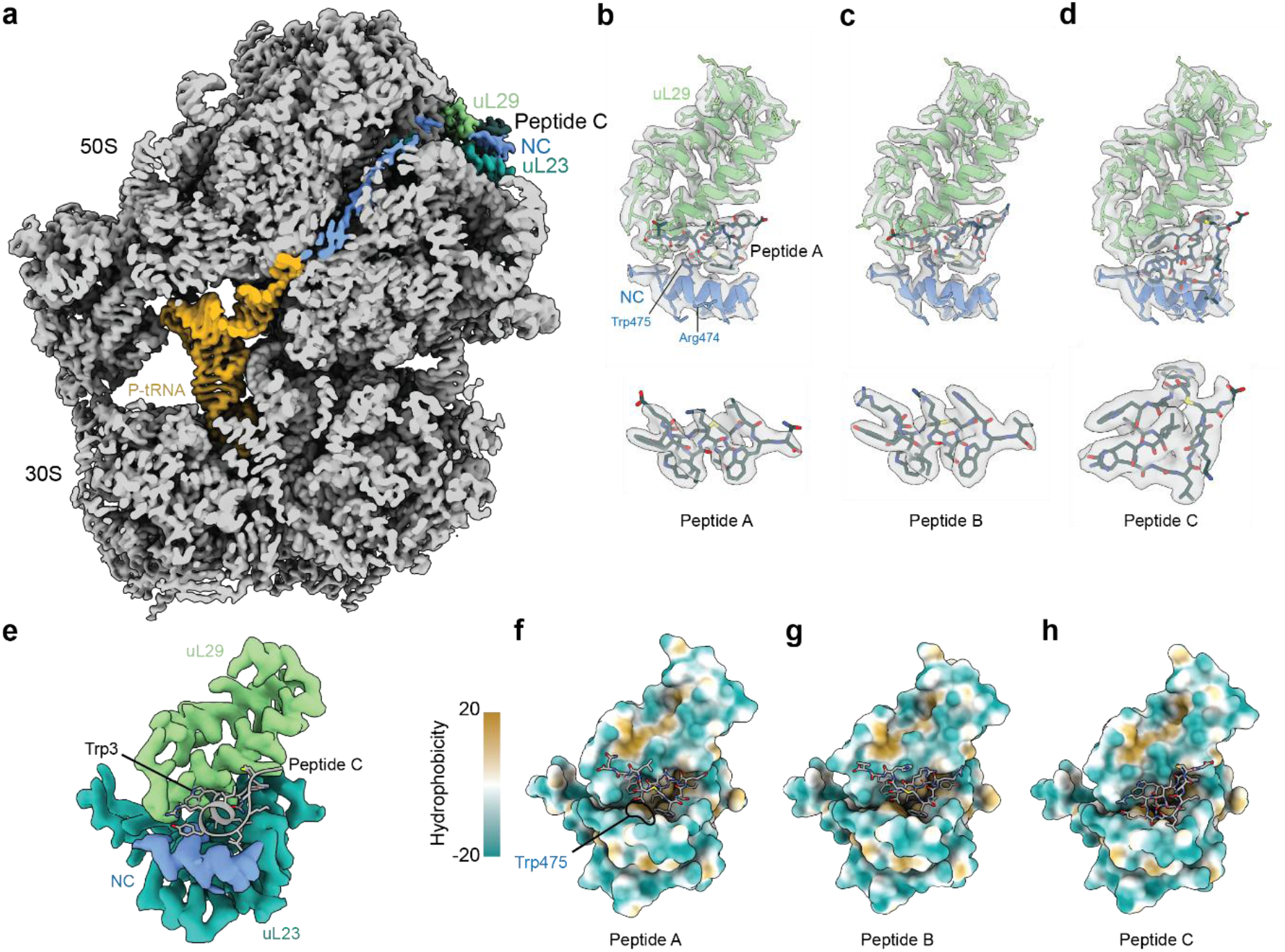
Cryo-EM reveals a composite peptide binding pocket on the ribosome surface. a, Overall structure of RNC510 bound to peptide C. Cross-section of cryoEM map showing the NC emerging from the ribosome exit port to interact with uL23, uL29 and peptide C. **b-d,** Peptide binding poses. CryoEM maps and atomic models are shown for each peptide in complex with uL29/NC (top) or alone (bottom). **e**, Peptides bind a pocket formed by uL23, uL29 and the NC. The atomic model of peptide C is shown together with the experimental density for the NC and ribosomal proteins. Trp3 of peptide C that interacts with uL29 is indicated. **f-g**, Surface representation of peptide binding pockets, colored by hydrophobicity.

The NC helix, uL23 and uL29 together create a hydrophobic cleft accommodating the cyclic part of each peptide (Fig. 5e-h). The contribution of the NC helix to this binding site explains why the peptides bind neither empty ribosomes nor ribosomes translating different sequences (Fig 2c-e). Similarly, the contribution of uL23 and uL29 to the binding pocket explains why the peptides do not bind full length β-gal or off-ribosome fragments of β-gal (Fig 2c-e). Our structures also explain why the peptides do not bind the 1-490 fragment of β-gal during ongoing translation (Fig. 3b-d). Although residues 466 to 478 are present in the 1-490 fragment, translation terminates with these residues still occluded within the exit tunnel, and thus unable to interact with uL23 and uL29 to form the peptide binding pocket.

In summary, NC interactions with the ribosome surface create a cryptic pocket comprising the NC and ribosomal proteins surrounding the exit port. This pocket is exploited by all three peptides to enable tight and selective binding to RNC_510_.

### A small structured element in the nascent chain suffices for high-affinity binding

We next probed the molecular determinants of peptide binding. Most contacts between the RNC and each peptide involve hydrophobic interactions (Fig. 6a and Extended Data Fig. 9a,b). We noted that the side chain of Trp475 from the NC is a prominent contributor to the overall hydrophobic cleft (Fig. 5f and 6a), and found that mutating it to alanine abolished the binding of all three peptides (Fig. 6b-d). By contrast, the adjacent Arg474 points away from the binding site and substituting it for alanine had no significant effect on binding (Fig. 6a-d). All three peptides contain a tryptophan residue which sits in a groove on the surface of uL29 (Trp10 in peptides A and B, Trp3 in peptide C), suggesting a key interaction for overall binding (Fig. 5e, 6a and Extended Data Fig. 9a,b). Indeed, mutating Trp10 to alanine in peptide A abolished binding (Fig. 6e). Similarly, Trp1 of peptides A and B occupies a pocket formed by uL29 and the NC, and mutating it to alanine in peptide A prevented binding (Fig. 6e).

**Fig. 6.**
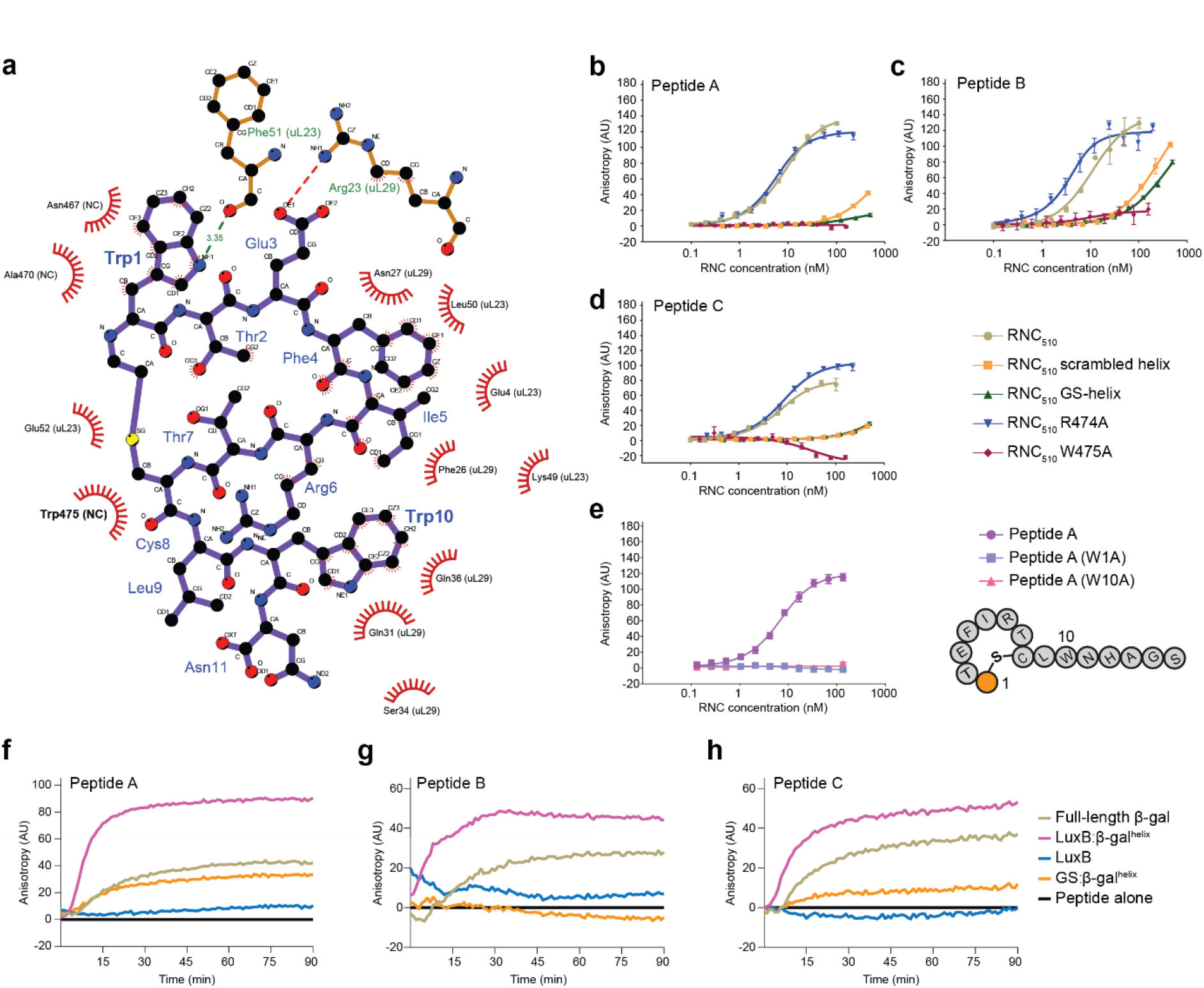
A small structured element in the nascent chain suffices for high-affinity binding. a, Map of interactions between peptide A and RNC510^38^. **b-d**, Peptide binding depends on hydrophobic contacts and folding of the NC helix. Binding to RNC510 with variant NCs was measured by fluorescence anisotropy as in Fig. 2. In wild-type RNC510 the helix-forming sequence is ^466^ANHDALYRWIKSV^478^. Scrambled helix: ^466^NADHARYLWIKSV^478^. GS-helix: ^466^ANGSAGSRGSKSV^478^. **e**, Hydrophobic residues in peptide A are required to bind RNC510. Binding was measured as in **b-d**. Data are presented as mean ± SD, n = 3 independent experiments. **f-h**, A short β-gal helix in the NC is sufficient for peptide binding to RNCs. Peptide binding during ongoing translation was measured as in Fig. 3, for chimeric constructs consisting of the β-gal helix and either LuxB (LuxB:β-gal^helix^) or a 100-aa GlySer-rich sequence (GS:β-gal^helix^).

Although the RNC contains 510 residues of nascent β-gal, our HDX-MS and cryoEM analysis suggested that only a short α-helical segment (466-478) in the NC contributes to peptide binding. Folding of the α-helix is necessary for binding to RNC_510_, as binding was abolished by either scrambling the helix-forming residues or introducing Gly-Ser substitutions (Fig. 6b-d). To test whether the β-gal helix is sufficient for peptide binding, we prepared a chimeric construct encoding residues 1-270 of the unrelated TIM-barrel protein, *Vibrio harveyi* LuxB, followed by residues 466-510 of β-gal (LuxB:β-gal^helix^). All 3 peptides bound LuxB:β-gal^helix^ during in vitro translation, but did not bind wild-type LuxB (Fig 6f-h). Moreover, the peptides bound a LuxB:β-gal^helix^ RNC with near-identical affinity to RNC_510_, indicating that the β-gal helix suffices to confer peptide binding (Extended Data Fig. 9c). To test whether binding is influenced by the overall foldedness of the NC, we replaced residues 1-465 of β-gal with a 100-amino acid GlySer-rich sequence, retaining the helix and C-terminal residues of β-gal (GS:β-gal^helix^). Peptides A and C bound GS:β-gal^helix^, demonstrating that binding can occur even if the NC is largely unstructured (Fig 6f-h). By contrast, peptide B did not bind GS:β-gal^helix^, possibly because the GlySer-rich sequence alters the stability or position of the NC helix.

Together, these results show that shape complementarity and interactions between hydrophobic residues drive cyclic peptide binding to the translation intermediate of β-gal. A short helical motif in the NC critically contributes to the binding pocket, and suffices for binding even in the context of an otherwise unstructured protein.

## Discussion

Here, we report a new class of binder that targets cotranslational folding intermediates, and describe a generalizable platform for their discovery (Fig. 7). Affinity panning against a stalled RNC identified cyclic peptides that bound tightly and specifically to a neo-interface formed by the NC and ribosome surface, even during real-time protein synthesis. The screening platform is agnostic to binder mechanism, and we anticipate that other binding modes can be exploited depending on the characteristics of the NC.

**Fig. 7.**
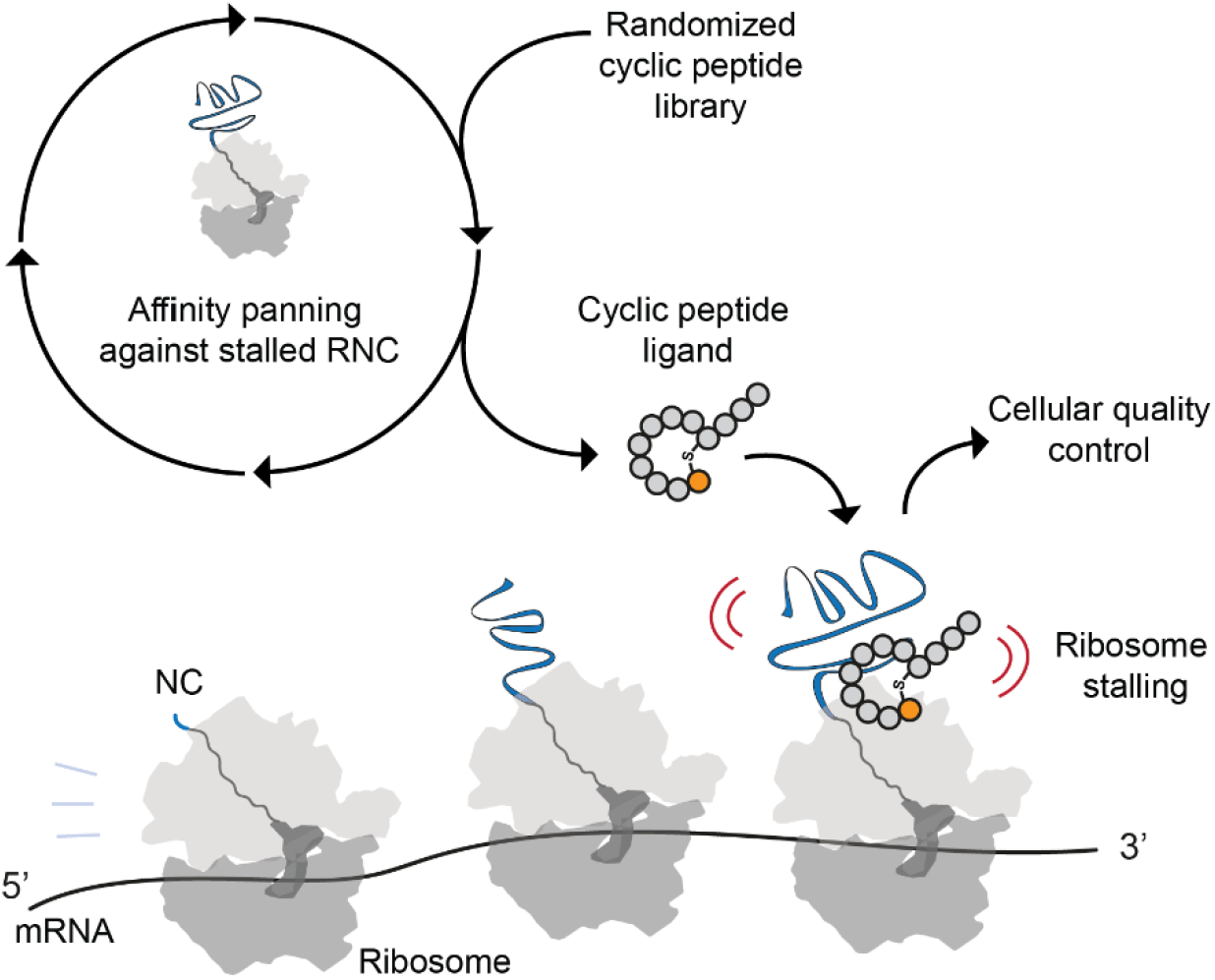
Towards selective drugs of protein biogenesis. Stalled RNCs can be used as a target for RaPID mRNA display, enabling discovery of cyclic peptides that bind specifically and with high affinity to folding intermediates transiently populated during protein biogenesis. A possible consequence of peptide binding is stalled translation, which in cells is expected to trigger quality control processes that disassemble the ribosome and degrade the NC.

Cotranslational binders may offer a new means to address challenging targets that do not expose easily druggable surfaces in their mature folded state, including intrinsically disordered proteins that are generally considered undruggable^39,40^. Intrinsically disordered proteins can transiently sample folded states, often α-helices, which become stabilized when bound to a partner^41–46^. As we show here, the ribosome surface can stabilize local NC structure to create a temporary binding pocket in an otherwise unstructured protein. Importantly, the fact that only a small region of the NC is involved in binding does not entail that binding is non-specific; point mutants or scrambled sequences are not recognized. This is because binding is determined not only by peptide:NC interactions, but also NC fold and NC:ribosome interactions, contrasting with approaches that target short linear sequence motifs deep in the ribosome exit tunnel^7^.

Although we show that the peptides bind in vitro, their selectivity in cells remains to be determined. *E. coli* is not an optimal model in this regard, because of the permeability barrier posed by the outer membrane^47^. Future efforts will focus on screening for binders of cotranslational protein folding intermediates in human cells, and test whether these binders exploit cellular quality control pathways that operate at the ribosome (Fig. 7). Ribosome processivity is closely monitored in human cells, with stalling (or even pausing) triggering quality control pathways that degrade the mRNA and NC^20^. Aberrant NCs are also ubiquitinated and degraded even in the absence of stalling^19,21–23^. Alternatively, peptide binders can be converted to PROTACs^3,48,49^, exploiting the exit vector at the site of mRNA attachment to the macrocycle. Cotranslational intermediates are likely to be particularly vulnerable to proximity-induced degradation because they are incompletely folded. Finally, since our screening approach does not depend on the consequence of binding, it may be possible to discover peptides that correct rather than disrupt folding.

## Acknowledgements

We thank the Crick Chemical Biology STP for peptide synthesis, Svend Kjaer for GFP-clamp beads, Simone Kunzelmann for help with optimising binding assays, Jaideep Jayachandran for help with photo-crosslinking, Sarah Caswell for advice guiding the project, and all members of Balchin and Walport labs for help and discussion. D.B. L.J.W and O.P. acknowledge funding from the Crick/AstraZeneca alliance (P2022-0010). D.B. and L.J.W. were also supported by the Francis Crick Institute which receives its core funding from Cancer Research UK (CC2025, CC2030), the UK Medical Research Council (CC2025, CC2030), and the Wellcome Trust (CC2025, CC2030). D.B acknowledges funding from UK Research and Innovation (UKRI) under the UK government’s Horizon Europe funding guarantee [FoldingMap, EP/X020428/1].

## Author contributions

O.P. performed all experiments except HDX-MS and Cryo-EM, and wrote the manuscript together with T.B.V, L.J.W. and D.B. T.B.V performed the cryo-EM experiments and data analysis. A.R. performed the HDX-MS experiments. C.M.J. designed and prepared the photo-Leu-containing peptide. J.S.D. contributed to peptide synthesis. A.N. collected cryo-EM data. R.D. contributed to initial project design. L.J.W. and D.B. conceived and supervised the project.

**Competing interests:** We declare no competing interests.

## Supplementary information

Supplementary Data S1. Sequences of top 1000 hit peptides from RaPID screen.

Supplementary Data S2. HDX-MS summary table and uptake data.

Extended Data Table 1. Sequences of purified peptides.

Extended Data Table 2. Cryo-EM data collection, map reconstruction and model refinement.

Extended Data Figures 1-9.

## Methods

### Purification of Ribosome Nascent Chain complexes (RNCs)

The design and purification of constructs encoding RNC_510_, RNC_GS_ and RNC_DHFR_ were described previously^10,11,26,35^. RNC purification was performed as previously described. In brief, E. coli BL21(DE3) cells were transformed through heat shock at 42 °C for 42 sec, with a plasmid encoding N-terminally GFP-tagged gene of interest with a C-terminal arrest-enhanced variant of the SecM ribosome arrest sequence^27^. Cells were grown in ZYM-5052 auto-induction media overnight at 37 °C and harvested by centrifugation at 4000 rpm for 20 min. The cell pellet was dissolved in RNC lysis buffer (70 mM Tris, pH 7.5, 150 mM KCl, 10 mM MgCl2, 8 U/mL RiboLock RNase inhibitor, 0.5x Halt Protease Inhibitor Cocktail (ThermoScientific)), snap-frozen in liquid nitrogen and stored at-80 °C. The cells were lysed by freeze thawing and lysozyme treatment (2 mg/mL lysozyme) for 15 min on ice. The lysate was further DNase treated (0.05 Kunitz units/μL RNase-free DNase (QIAGEN)) and clarified by centrifugation at 13000 rpm for 20 min in 4 °C. The clarified lysate was layered onto a 35% sucrose cushion (35% sucrose, 50 mM HEPES-NaOH pH 7.5, 12 mM Mg(OAc)2, 1 M KOAc, 1 mM DTT, 8 U/mL RiboLock RNase inhibitor and 0.2x Halt Protease Inhibitor Cocktail) to enrich for ribosomes ultracentrifugation. Layered samples were centrifuged at 70,000 rpm in a Beckmann TLA-110 rotor for 2 h at 4 °C. Supernatants were discarded and the pellets containing ribosomes and RNCs were resuspended with gentle agitation overnight in RNC buffer (50 mM HEPES-NaOH pH 7.5, 12 mM Mg(OAc)2, 100 mM KOAc, 1 mM DTT and 8 U/mL RiboLock RNase inhibitor) at 4 °C. Resuspended ribosomal pellets were incubated with RNC buffer-equilibrated in-house-prepared GFP-clamp coupled agarose beads overnight at 4 °C on a spinning wheel (300 μL of beads for 200 mL of culture)^50^. These beads with immobilized RNC were used for RaPID screening as described below.

For biophysical characterization and cryoEM structure determination, RNCs were eluted from the beads by cleaving from the GFP tag with 3C protease, overnight at 4 °C. The eluted RNC was purified by second round of ultracentrifugation as above, the pellet was washed and resuspended in RNC buffer, snap-frozen in liquid nitrogen and stored at-80 °C. The RNC concentration was determined by measuring A_260_.

### RaPID screening

A standard RaPID protocol was followed as previously described^51,52^. In brief, the selection process began with preparation of a semi-randomized mRNA library consisting of the sequence AUG-(NNK)_6-12_-UGC. This library was conjugated to puromycin bearing DNA linker at 1:1 ratio. Following ligation reaction, the library was purified by precipitation and resuspended in Milli-Q water. This library was then introduced to a Flexizyme (flexible tRNA-acylation ribozymes) mediated Flexible In vitro Translation (FIT) system^53,54^. Here, the initiation AUG codon was reprogrammed to encode the non-canonical N-chloroacetyl-D-Tryptophan (ClAc-D-Trp) in a methionine (Met)-depleted PURE system. For this, tRNA^ini^_CAU_ was aminoacylated using the cyanomethyl ester of ClAc-D-Trp with flexizyme, eFx. Following translation, libraries were treated with EDTA (37.5 mM) to dissociate the ribosome complex and allow spontaneous peptide cyclisation between the N-terminal chloroacetyl group and a cysteine thiol, yielding macrocyclic peptides with non-reducible thioether linkages. The cyclized peptide-mRNA library was then reverse-transcribed and desalted into selection buffer (RNC buffer with 0.1% Tween20) before dilution with equal volume of 2X blocking solution (selection buffer containing 0.2% (w/v) acylated bovine serum albumin (BSA) and 8 mg/mL yeast tRNA). The first round of selection was performed at a 50 μL translation scale. 200 nM of bead bound RNC_510_ was washed thrice with selection buffer (50 μL, 9000 rpm, 1 min) and then incubated with 1X blocking solution for 30 mins, followed by another 3 washes with selection buffer (50 μL, 9000 rpm, 1 min). The desalted peptide library mix was then added to bead bound RNC_510_ and incubated for 30 min by gentle agitation at 4 °C. The sample was washed thrice with selection buffer to remove non-binders (100 μL, 9000 rpm, 1 min) and the bound library was recovered by heat induced elution and PCR amplification.

Negative selection was omitted in Round 1 to maximise library diversity. Subsequent rounds were performed on a 5 μL scale and from Round 2 onward, the amplified library was pre-incubated with 200 nM bead bound RNC_GS_ (negative selection) for 30 mins to deplete non-specific binders. The resulting non-binders were then transferred to the target RNC for positive selection. The affinity panning was terminated after round 6. DNA pools from both positive and negative selection steps across all rounds from round 2 onward were analysed by Next-Generation Sequencing (NGS) to monitor the convergence of the library and identify highly enriched peptide motifs. The resulting base calls were parsed to obtain the DNA sequences, which were subsequently translated in silico to peptide sequences and ranked by NGS sequence enrichment score. The top 1000 sequences from the final positive selection (round 6) were clustered using hierarchical agglomerative clustering with Ward linkage in Euclidean space using a custom python script (Data S1)^33,34^.

### Peptide synthesis

All the peptides used in this study were synthesised by solid phase peptide synthesis (SPPS) using fluorenylmethyloxycarbonyl (fmoc) chemistry and an automated synthesiser, Prelude X (Gyros Protein Technologies) (Extended Data Table 2). Peptides were synthesized as amides using a low load Rink Amide resin (0.29 mmol/g, Novabiochem). Each amino acid was coupled with 5 eq. Fmoc-protected amino acid, 5 eq. N,N-diisopropylcarbodiimide (DIC), 5 eq. Oxyma pure in dimethylformamide (DMF). After each coupling, capping was carried out with 20 eq. acetic anhydride and 20 eq. N,N-diisopropylethylamine (DIPEA) in DMF. The following fmoc-deprotection was performed with 20% piperidine, 1% formic acid in DMF. After deprotection of the last amino acid, 5 eq. N-(chloroacetoxy)-succinimide in DMF was coupled to the N-terminus (2 h, RT). Peptides were deprotected and cleaved off the resin with trifluoracetic acid (TFA): triisopropyl silane (TIPS): 2,2-(ethylenedioxy)-diethanethiol (DODT): water (91:3:3:3) (2h, RT), except peptide D1L10PhoL which was cleaved with TFA:TIPS:water:thioanisole (90:5:2.5:2.5) (4 h, RT). Crude peptides were precipitated in cold diethyl ether, washed with diethyl ether and dissolved in dimethylsulfoxide (DMSO):water (80:20). The peptide mixtures were mixed overnight at room temperature to allow for decarboxylation. Next day, triethylamine was added until pH 9 for cyclisation. The crude peptides were purified with reverse phase HPLC and lyophilised. The purified peptides were analysed with HPLC-MS (Extended Data Figure 2). The purified cyclic peptides were analysed on a Waters HPLC system (2489 UV/VIS detector) linked with a MS detector (SQ detector 2) with a C18 column (XBridge BEH C18 column, 4.6 x250 mm, 5 µM). Solvent A: 0.1% TFA in water, Solvent B: 0.1 % TFA in acetonitrile. Analytical HPLC Method: 10% B 0-2.5 min, 10-90% B 2.5-20 min, 90-98% B 20-22 min, 98% B 22-27 min, 10% B 27-32 min.

### FAM conjugation

The peptides were dissolved in DMSO and 2 equivalents of FAM DBCO (Lumiprobe) were added. After the sterically promoted cycloaddition reaction between DBCO and the azido-group on the C-terminal lysine had gone to completion (overnight incubation at room temperature), the peptides were purified and analysed as described above (Extended Data Table 2 and Extended Data Figure 2).

### Fluorescence anisotropy

The FAM-conjugated peptides were dissolved in DMSO and diluted using RNC buffer to a final peptide concentration of 5 nM (for binding to RNCs) or 25 nM (for binding to soluble full length β-gal) and final DMSO < 0.5 % v/v. Peptides were incubated with a 12-point serial dilution of target RNC or protein (ranging from 0 to 1000 nM) for 10 min at 4 °C. Fluorescence anisotropy was measured using a CLARIOstar plate reader. Following baseline subtraction, the anisotropy values were used to generate binding isotherms. Equilibrium dissociation constants (*K*_D_) of binding were determined by quadratic fits to the data using GraphPad Prism. Experiments were performed in triplicate and used to calculate the reported mean.

### Coupled Fluorescence anisotropy / in vitro transcription/translation (IVTT)

IVTT reactions were performed using the PURExpress In Vitro Protein Synthesis Kit (NEB) as per the manufacturer’s instructions. Solution A (3 µL), Solution B (4 µL), and RNase inhibitor (0.2 µL, RNasin Ribonuclease Inhibitor, Promega) were mixed on ice with FAM-conjugated peptide and the template plasmid at a final concentration of 10 nM and 12 ng/µL respectively. A peptide-only reaction was also performed to determine the baseline. The IVTT reaction was initiated by incubating the plate at 37 °C within a CLARIOstar plate reader, with fluorescence anisotropy recorded every 60 sec for 90 min. Following baseline subtraction, the anisotropy value was used to generate kinetic profiles using GraphPad Prism. Experiments were performed in triplicate and used to calculate the reported mean.

### Photo-crosslinking

A photo-activatable derivative of Peptide C was synthesized, in which the leucine residue at position 10 was substituted with L-photoleucine. This derivative was then conjugated with FAM for detection. Peptide C L10PhoL is crosslink-active upon UV irradiation at 365 nm. The peptide was added to IVTT reaction components (3 µL Solution A, 4 µL Solution B, and 0.2 µL RNasin Ribonuclease Inhibitor, Promega) and template DNA (12 ng/µL) in PCR tubes. The IVT reaction was initiated by incubating the tubes in a thermomixer preheated to 37 °C with shaking at 350 rpm. After 10 min, the standard lid was replaced with a custom-built lid equipped with a UV irradiation source for photo-crosslinking. The reaction was allowed to proceed under continuous UV (365 nm) exposure for an additional 10 min before quenching by addition of an equal volume of 4X Laemmli sample buffer and incubation at 60 °C for 5 min. The samples were resolved via 4-12% SDS-PAGE and in-gel FAM fluorescence was recorded using a Typhoon FLA 9500 biomolecular imager. The gel was then stained with Coomassie to visualize overall protein production.

### HDX-MS

HDX-MS was conducted as described previously^26^ with some modifications. In short, stock samples of empty ribosomes (NEB, P0763S), RNC_1-510_, and RNC_1-510_ mixed with 16 μM of individual peptides, were prepared in RNC low-salt buffer at final 5.5 μM ribosome concentration. Stock of full-length β-galactosidase was prepared in RNC low salt buffer at final 15 μM concentration. All stocks were subsequently incubated for 30 min at 25 °C.

Deuterium labelling of RNC-containing samples was initiated by mixing 3 μL of stock samples with 27 μL deuteration buffer (10 mM HEPES-NaOH, pH 7.1, 30 mM KOAc, 12 mM Mg(OAc)2, 1 mM DTT, RiboLock RNase inhibitor (ThermoScientific), 97% D2O) at 25 °C. After labelling at 25 °C for 10, 100 or 3000 s, the reaction was quenched with an equal volume (30 μL) of ice-cold quench buffer (100 mM sodium phosphate, pH 1.4, 4 M guanidium hydrochloride, 10 mM TCEP). The pH after quenching was 2.5. Digestion was initiated by addition of 20 μL pepsin-agarose 50% v/v bead slurry (prepared in-house as described previously^26^) equilibrated in 0.1% formic acid. Following 100 s of digestion at 10 °C with rapid mixing every 30 seconds, the sample was centrifuged (13,000 g, 15 s, 0 °C) through 0.22 μm PVDF filters (Millipore, UFC30GVNB). The flow through was immediately snap-frozen in liquid nitrogen for short-term storage. The same protocol was followed to prepare undeuterated controls, except the deuteration buffer was replaced by a H-based buffer (10 mM HEPES-NaOH in H2O, pH 7.5, 30 mM KOAc, 12 mM Mg(OAc)2, 1 mM DTT, RiboLock RNase inhibitor).

Frozen samples were rapidly thawed and immediately injected into an Acquity UPLC M-class system with the cooling chamber containing the chromatographic columns kept at 0 ± 0.2 °C throughout data collection. Peptic peptides were trapped and desalted for 4 min (200 μL/min) on a 2.1 mm X 5 mm, C4 trap column (Acquity BEH C4 Van-guard pre-column, 1.7 μm, Waters, 186004623) then separated on a reverse phase Acquity UPLC HSS T3 column (1.8 μm, 1 mm X 50 mm, Waters, 186003535) at a flow rate of 90 μL/min. Peptides were eluted over 25 min using a 3-30% gradient of acetonitrile in 0.1% formic acid. Analysis was performed using a Waters Synapt G2Si HDMSE instrument in ion mobility mode, acquiring in positive ion mode over a range of 50 to 2,000 m/z with the conventional electrospray ionisation source. Calibration of the mass spectrometer was achieved using [Glu1]-Fibrinopeptide B (50 fmol/μL, Merck, F3261) and the instrument was operated at a source temperature of 80 °C with the capillary set to 3 kV.

MSE data were processed using Protein Lynx Global Server (PLGS, Waters) to identify peptides in the undeuterated control samples using information from a non-specific cleavage of a database containing sequences of E. coli β-galactosidase, 70S ribosomal proteins as well as porcine pepsin. PLGS search was performed using energy thresholds of low = 135 counts and elevated = 30 counts. Peptides identified by PLGS were subsequently filtered and processed in DynamX (Waters) with filters of minimum products per amino acid of 0.05 and minimum consecutive products of 1. All spectra were manually inspected, and poor-quality assignments were removed. Any peptides assigned to nascent chains that were also detected in empty ribosome control samples were removed. Additionally, peptides assigned to β-galactosidase in nascent chains but not in isolated β-galactosidase, were also removed. Relative deuterium uptake in Da was calculated by subtracting the centroid mass of undeuterated peptides from those of deuterated peptides. Determined mean values of deuterium uptake are relative as they are not corrected for back-exchange. All HDX-MS experiments were performed in technical duplicates or triplicates for each RNC sample at each deuteration time and used to calculate the reported mean. All plotted uptake data, peptide coverage maps, number of replicates per sample and a summary of experimental conditions^55^ are shown in Data S2. The MS data have been deposited to the ProteomeXchange Consortium via the PRIDE partner repository^56^ with the dataset identifier PXD082965.

## CryoEM

### CryoEM grid preparation and data collection

RNC samples were diluted to 0.8 µM and incubated with 16 µM of each peptide (individually) at 25 °C for 30 min. Quantifoil R1.2/1.3 Cu-400 grids (Agar Scientific) were glow-discharged in air for 30 s with a GloQube instrument (Quorum Technologies). 4.5 µl of sample was applied to the grids before blotting with a Vitrobot Mark IV (Thermo Fisher Scientific) at 22 °C and 80% humidity with a blot time of 3 s and blot force of-1. Grids were plunge-frozen in liquid ethane and stored in liquid nitrogen.

Movies were collected on a 300-kV Titan Krios EM instrument (Thermo Fisher Scientific) with a Falcon 4i direct electron detector (Thermo Fisher Scientific) and a Selectris energy filter (Thermo Fisher Scientific) operating at a slit width of 10 eV. For each dataset, 10,000 movies were collected with EPU version 3.8.1.7603 using aberration-free image shift and fringe-free illumination at a nominal magnification of ×130,000 for a pixel size of 0.95 Å per pixel. Data collection details are in Extended Data Table 2.

### Data processing

Detailed image processing workflows are presented in Extended Data Figures 4-6. All three datasets were pre-processed in Relion 4^57^ using MotionCor2^58^ and CTFFIND4^59^ for motion correction and CTF estimation, respectively. Each dataset was processed separately.

For the datasets corresponding to peptides A and B, pre-processed micrographs were imported into Cryosparc v4^60^ and subjected to blob picking (particle diameter between 180 Å and 250 Å). Particles were extracted (binned 4 times) and cleaned up with three rounds of 2D classification. Particles from well-resolved 2D classes were selected for *ab initio* reconstruction (1 class) and homogeneous refinement. Focused classification with a mask around the tRNA binding sites (6 classes, 12 Å filter resolution) was performed and classes containing tRNA density in all three sites (A, P and E) were selected for reextraction (first binned 2 times, then unbinned) and homogeneous refinement. 3D classification (4 classes, 3 Å filter resolution) was performed to remove particles limiting the resolution and all good particles were taken to Relion 4 for 3D refinement, CTF refinement and Bayesian polishing.

For the dataset corresponding to peptide C, all image processing was performed in Relion 4 following a similar approach. Particles were autopicked using a Laplacian-of-Gaussian filter (particle diameter between 180 Å and 250 Å). Particles were extracted (binned 4 times) and cleaned up with three rounds of 2D classification. Particles from well-resolved 2D classes were selected to generate an initial model before 3D refinement. Focused classification with a mask around the tRNA binding sites (5 classes, 25 iterations, T=4) was performed and classes containing tRNA density in all three sites (A, P and E) were selected for reextraction (first binned 2 times, then unbinned) and 3D refinement before CTF refinement and Bayesian polishing.

### Model building and refinement

For each map, the structure of the apo RNC_510_ (PDB 9R3A^11^) was rigid-body fitted in ChimeraX v1.9^61^. The peptides were built in WinCoot v0.9^62^ with a glycine in position 0 to mimic the covalent bond between the first residue and the cyclising cysteine. The resulting models were refined with Isolde^63^ in ChimeraX, with a position restraint of 1.8 Å and 1,000 N between the α-carbon of Gly0 and the sulfur of the cyclising cysteine. The A-site tRNA was mutated in ChimeraX to reflect the tRNA most likely to recognise the CCC codon (ProL) at this position of the stalling sequence. The structure of the *E. coli* Gly-tRNA was predicted with AlphaFold3^64^ and fitted with Isolde into the E-site density. As the *E. coli* genome encodes 3 Pro-tRNAs able to recognise the CCU codon and occupy the P-site of our construct^65^, all three structures were predicted with AlphaFold3 and flexibly fitted into the density with Isolde before assessing the map-model agreement with per-residue Q-scores^66^ and visual inspection. This revealed that the ProM tRNA best matched our data, and was thus used for model building. Water molecules were added with the ‘Find Waters’ tool in WinCoot (peaks above 1.4 rmsd, minimum distance of 2.4 Å, maximum distance of 3.2 Å) before automatically deleting all water molecules which matched one or more of the following criteria: B-factor over 20 Å^2^; map rmsd level below 1 e^-^/Å^2^; closest contact less than 2.3 Å away; closest contact more than 3.5 Å away. Finally, each remaining water molecule was checked manually and removed or adjusted when needed. Each model was refined against its corresponding consensus map with phenix.real_space_refine in PHENIX version 2.0^67^ (8 macrocycles) after truncating the carbonyl oxygen from Gly0 and adding a covalent bond between the α-carbon of Gly0 and the sulfur of the cyclising cysteine. The refinement parameters included the covalent bond between the P-tRNA and the nascent chain (ideal distance of 1.4 Å) and the cyclisation bond (ideal distance of 1.8 Å). Model validation was performed in PHENIX (Extended Data Table 2).

## Data availability

Sequencing data from RaPID have been deposited in the Sequence Read Archive with the accession PRJNA1537536.

All mass spectrometry data have been deposited to the ProteomeXchange Consortium via the PRIDE partner repository^56^ with the dataset identifier PRIDE: PXD082965.

The cryoEM maps have been deposited in the Electron Microscopy Data Bank (EMDB), and the coordinates have been deposited in the PDB under the following accession numbers:

• Peptide A: PDB 34IA, EMDB 59858

• Peptide B: PDB 34XA, EMDB 100214

• Peptide C: PDB 34XQ, EMDB 100227

## Extended Data

**Extended Data Fig. 1.**
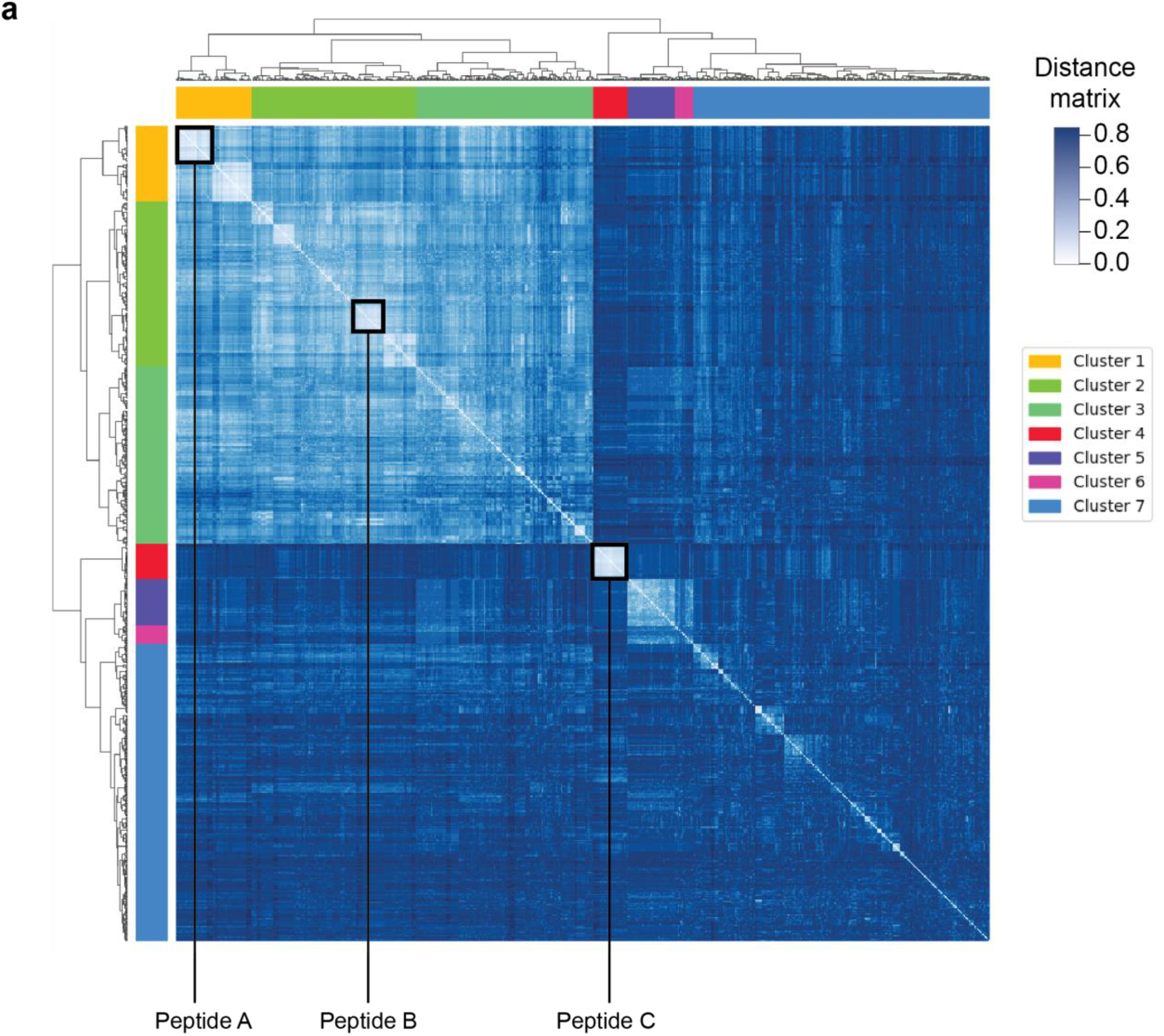
Clustering of sequences encoding hit peptides. a, Sequence clustering of the top 1000 most enriched peptides from RaPID screening. Lighter colors in the distance matrix indicate higher sequence similarity.

**Extended Data Fig. 2.**
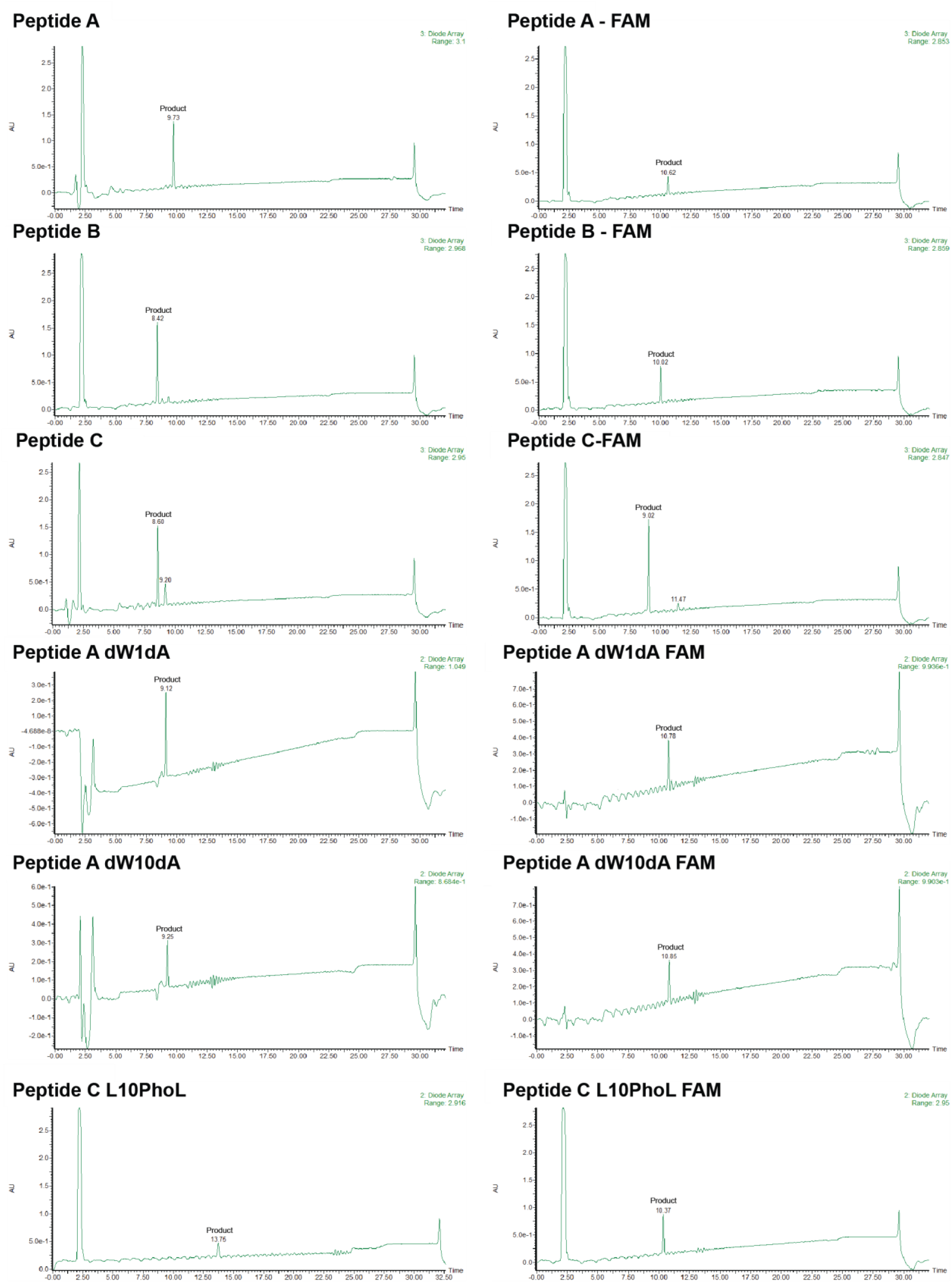
**HPLC analysis of purified peptides.**

**Extended Data Fig. 3.**
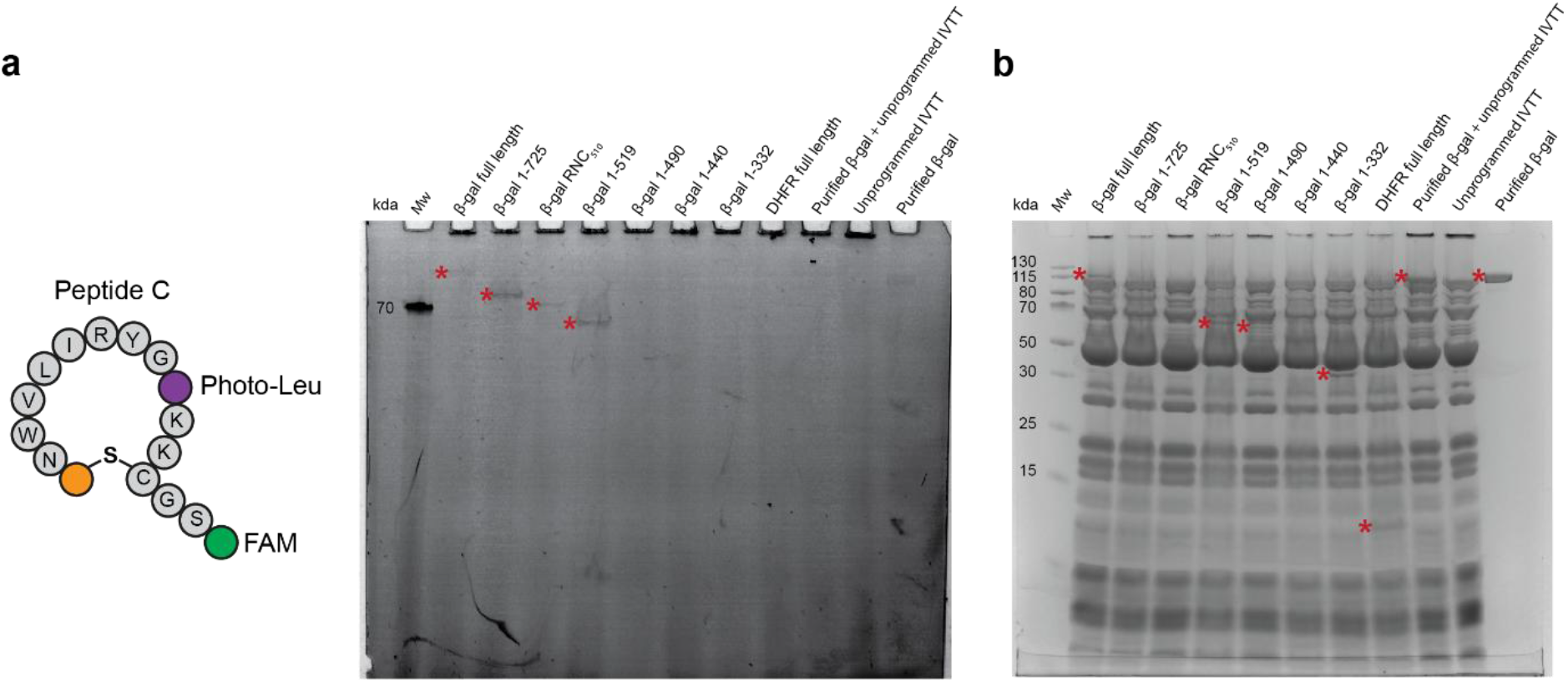
Peptide C specifically crosslinks to in vitro translated β-gal. IVTs were programmed with plasmids encoding DHFR or different lengths of β-gal, as in Fig. 3, in the presence of a Photo-Leu-containing variant of Peptide C labelled with FAM at the C-terminus. Crosslinking was induced by UV exposure and the reactions were resolved by SDS-PAGE. Crosslinked products were detected by FAM fluorescence (**a**), and newly-synthesized protein was detected by Coomassie staining (**b**). Where visible, crosslinked products and newly-synthesized protein are marked with an asterisk.

**Extended Data Fig. 4.**
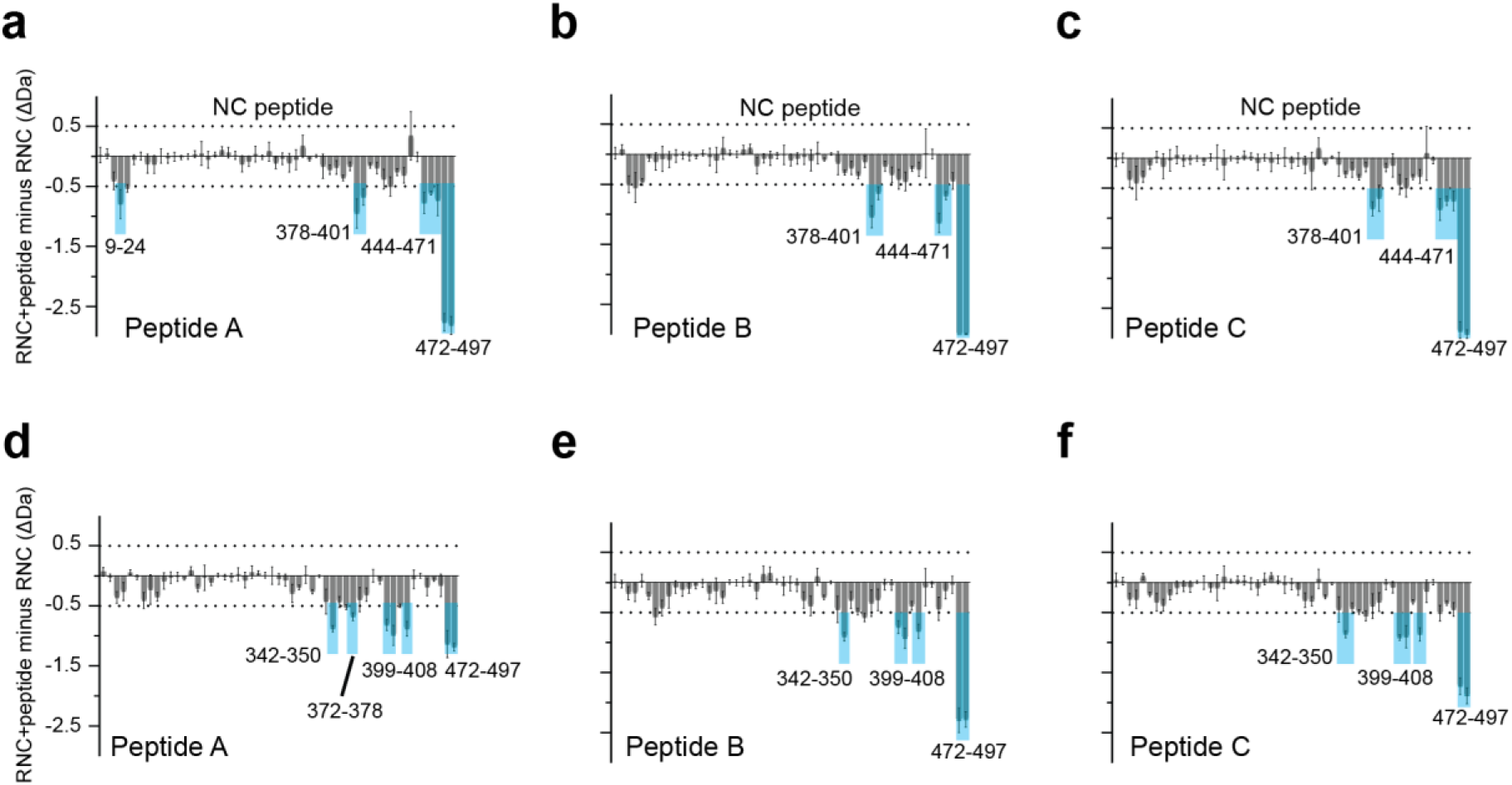
HDX-MS analysis of peptide binding to the NC in RNC510. Protection of β-gal NC peptides induced by cyclic peptide binding. Difference in deuteron uptake after 100 s (**a-c**) or 3000 s (**d-f**) between RNC510 alone and in complex with each cyclic peptide. Negative values indicate protection from exchange in the presence of the peptide. Peptides showing uptake differences ≥0.5 Da are labelled. HDX-MS data are presented as mean ± SD, n = 3 independent labelling reactions.

**Extended Data Fig. 5.**
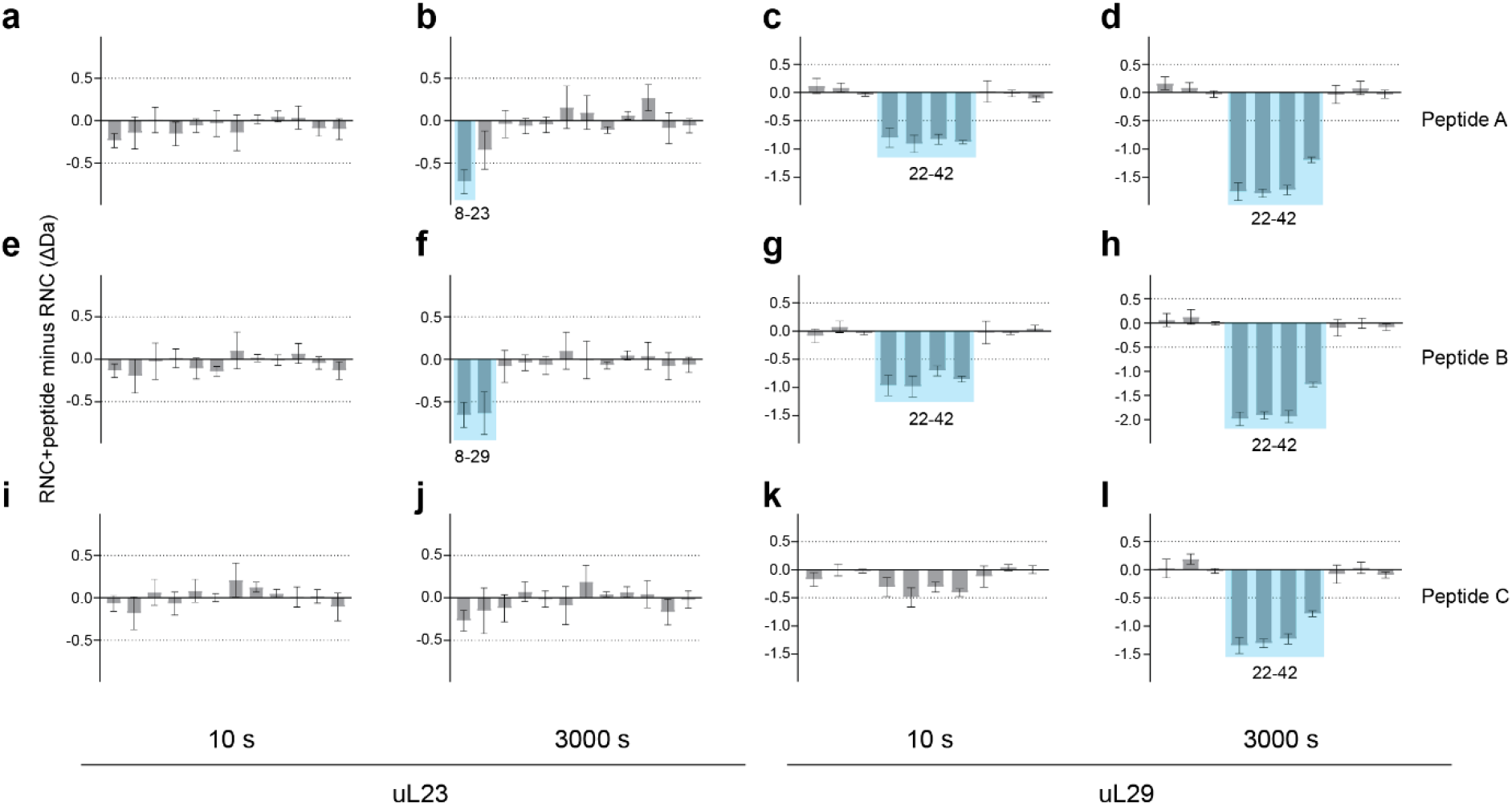
HDX-MS analysis of peptide binding to uL23 and uL29 in RNC510. Protection of ribosomal protein peptides induced by cyclic peptide binding. **a-l**, Difference in deuteron uptake after 10 or 3000 s, between RNC510 alone and in complex with each cyclic peptide. Negative values indicate protection from exchange in the presence of the peptide. Peptides showing uptake differences ≥0.5 Da are labelled. HDX-MS data are presented as mean ± SD, n = 3 independent labelling reactions.

**Extended Data Fig. 6.**
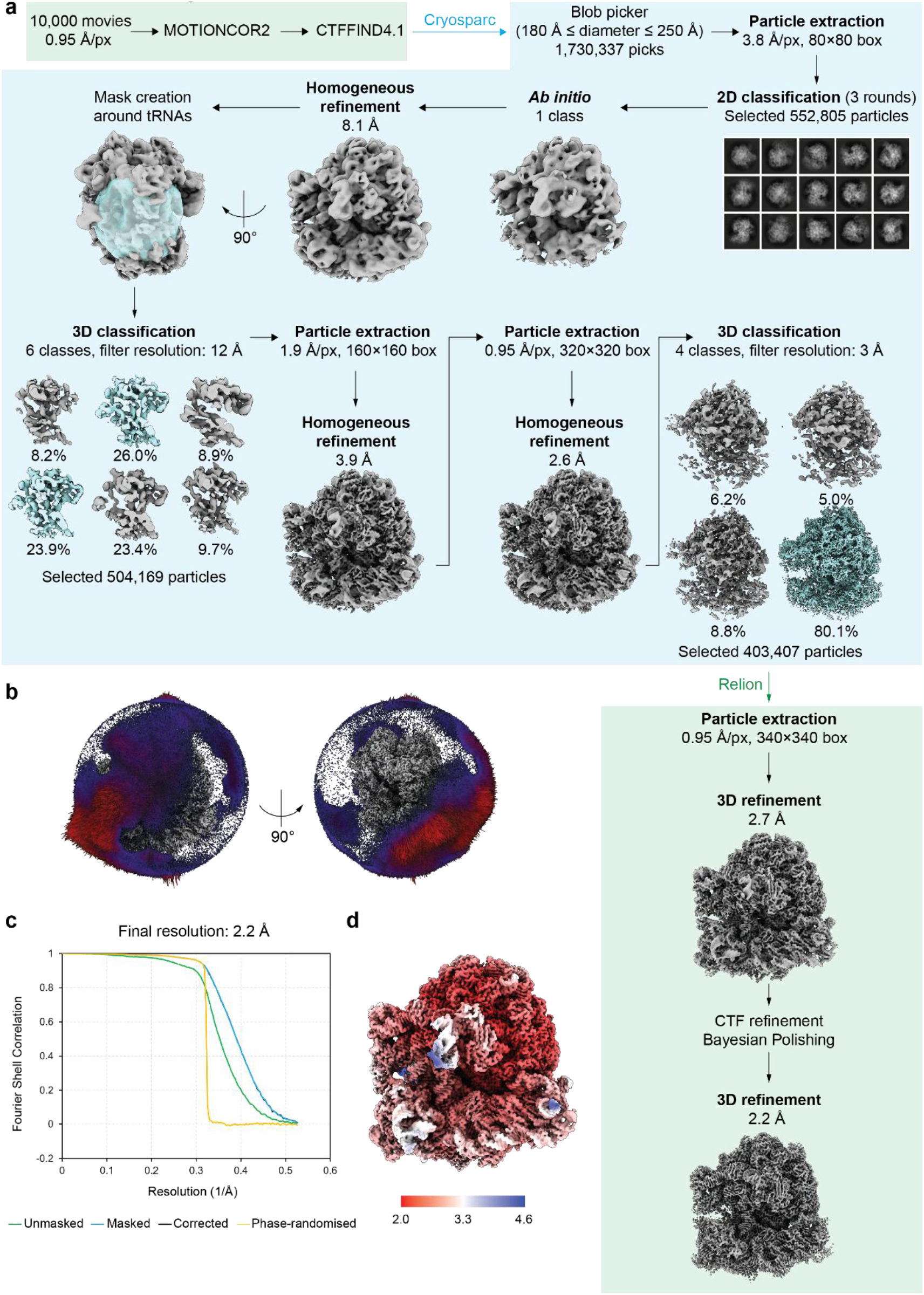
Cryo-EM data processing workflow for the complex between RNC510 and peptide A. a, Image processing workflow. **b**, Angular orientation distribution. **c**, Fourier Shell Correlation (FSC) curves of the half-maps. The FSC cut-off for global resolution measurement is 0.143. **d**, Local resolution of the consensus map, ranging from 2.0 Å (red) to 4.6 Å (blue).

**Extended Data Fig. 7.**
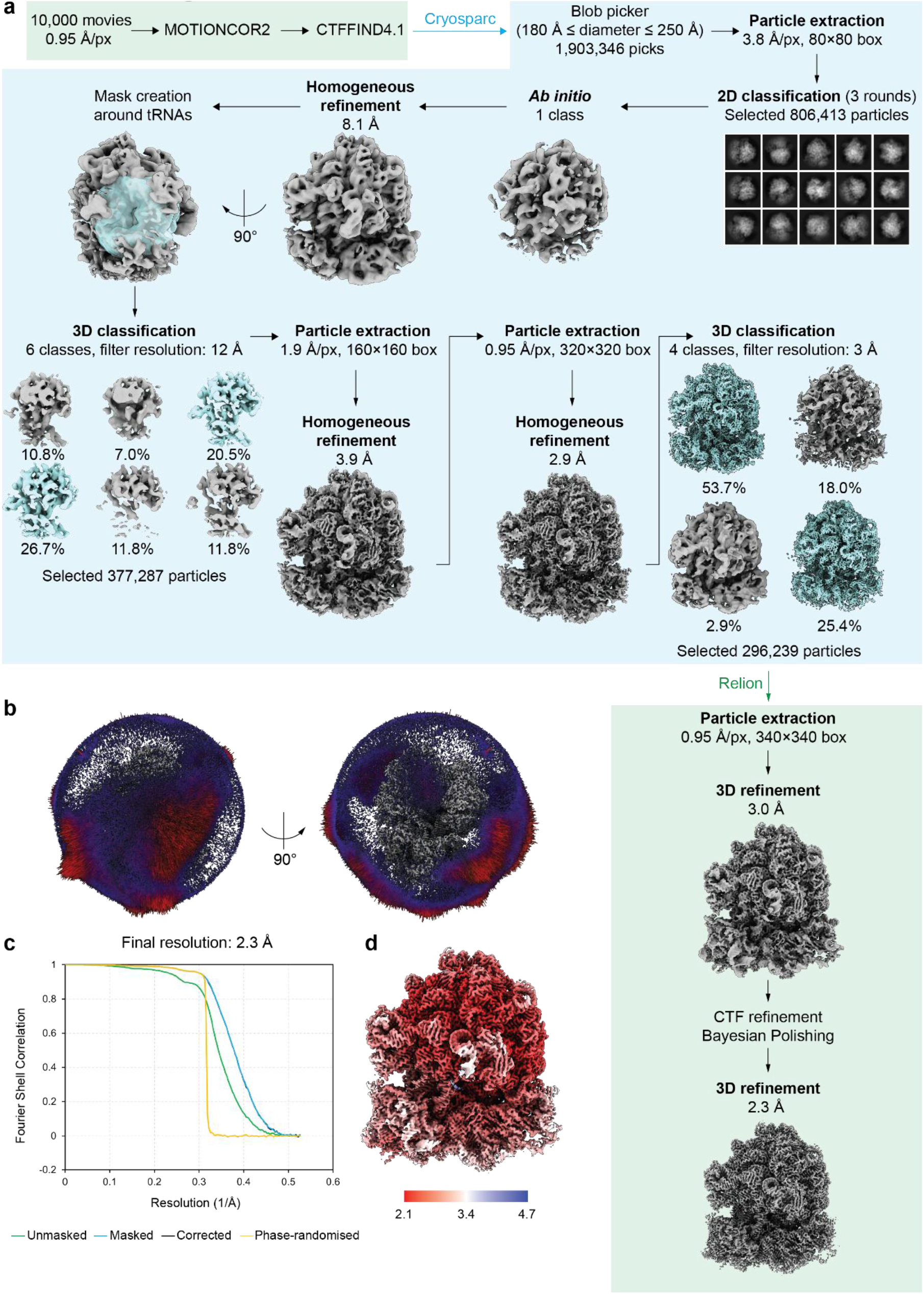
Cryo-EM data processing workflow for the complex between RNC510 and peptide B. a, Image processing workflow. **b**, Angular orientation distribution. **c**, Fourier Shell Correlation (FSC) curves of the half-maps. The FSC cut-off for global resolution measurement is 0.143. **d**, Local resolution of the consensus map, ranging from 2.1 Å (red) to 4.7 Å (blue).

**Extended Data Fig. 8.**
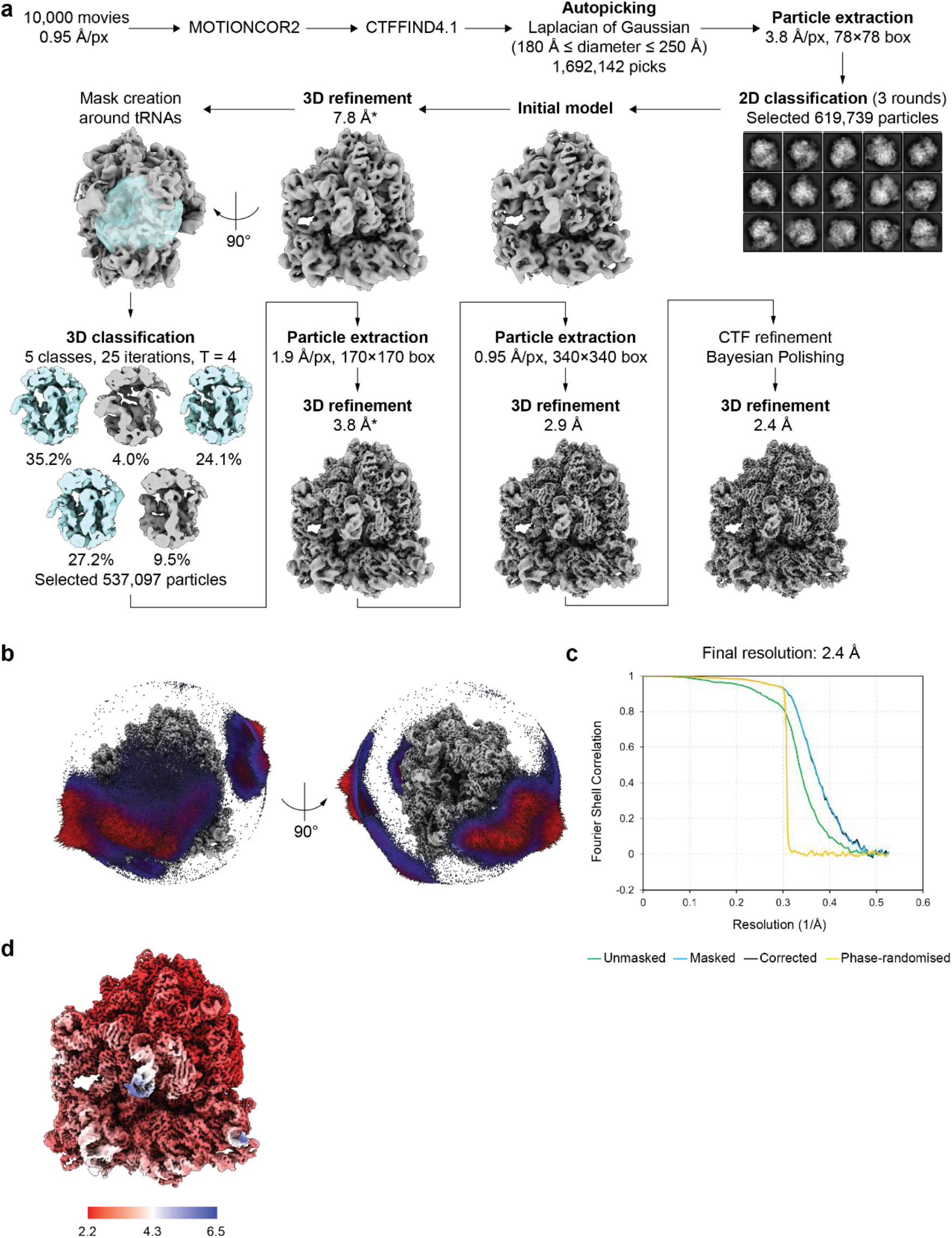
Cryo-EM data processing workflow for the complex between RNC510 and peptide C. a, Image processing workflow. *Unmasked resolution estimates. **b**, Angular orientation distribution. **c**, Fourier Shell Correlation (FSC) curves of the half-maps. The FSC cut-off for global resolution measurement is 0.143. **d**, Local resolution of the consensus map, ranging from 2.2 Å (red) to 6.5 Å (blue).

**Extended Data Fig. 9.**
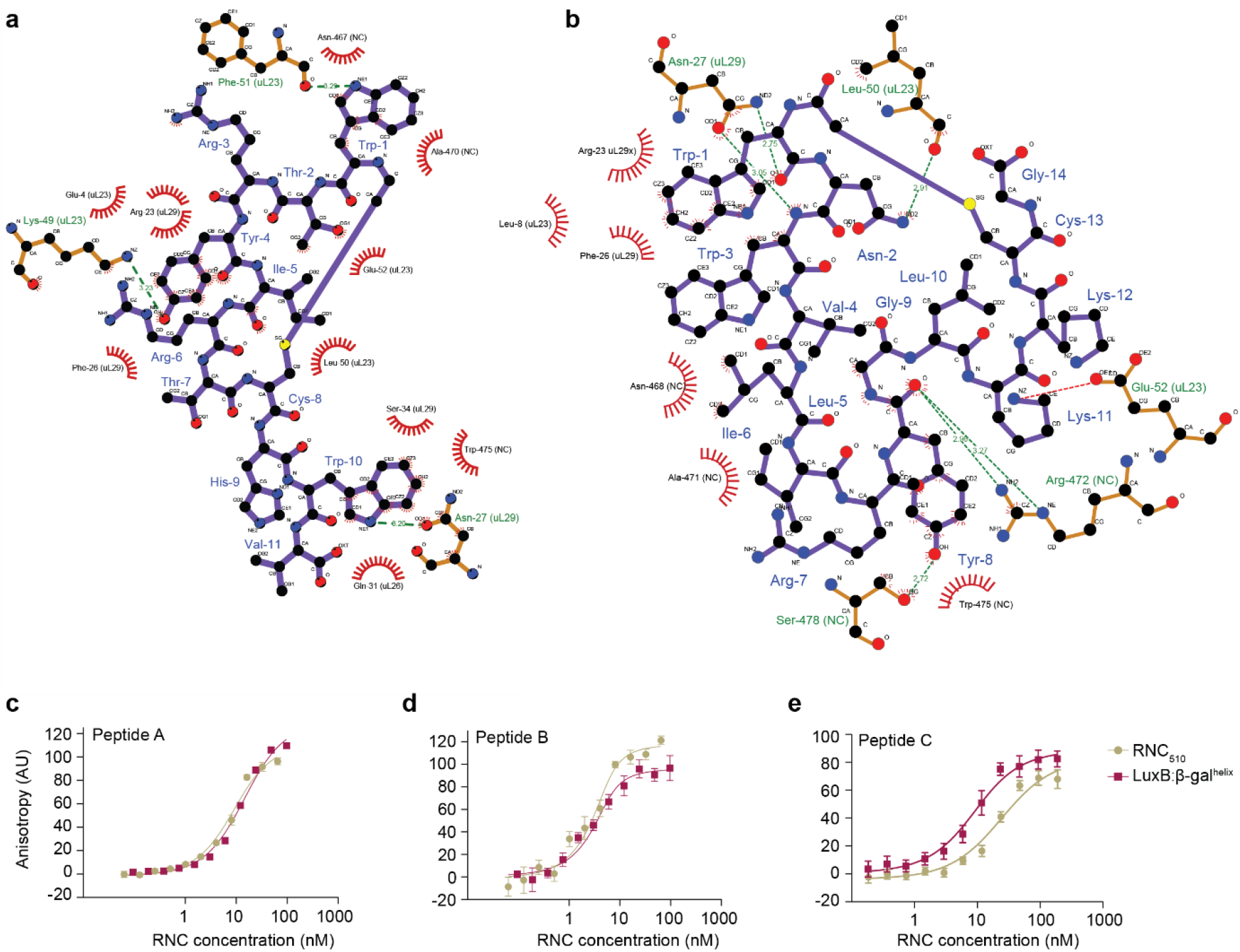
Molecular determinants of peptide binding to RNC510. a, Map of interactions between RNC510 and peptide B^38^. **b**, Map of interactions between RNC510 and peptide C. **c-e**, Quantification of peptide binding. Carboxyfluorescein-labelled peptide was incubated with increasing concentrations of different RNCs and binding was measured by the increase in fluorescence anisotropy. Data are presented as mean ± SD, n = 3 independent experiments. Dissociation constants are as follows. Peptide A: 7 ± 2 (RNC510) and 12.0 ± 0.2 (LuxB:β-gal^helix^); Peptide B: 0.8 ± 0.1 (RNC510) and 0.9 ± 0.6 (LuxB:β-gal^helix^), Peptide C: 7 ± 1 (RNC510) and 22 ± 3 (LuxB:β-gal^helix^).

---

**Extended Data Table 1.**
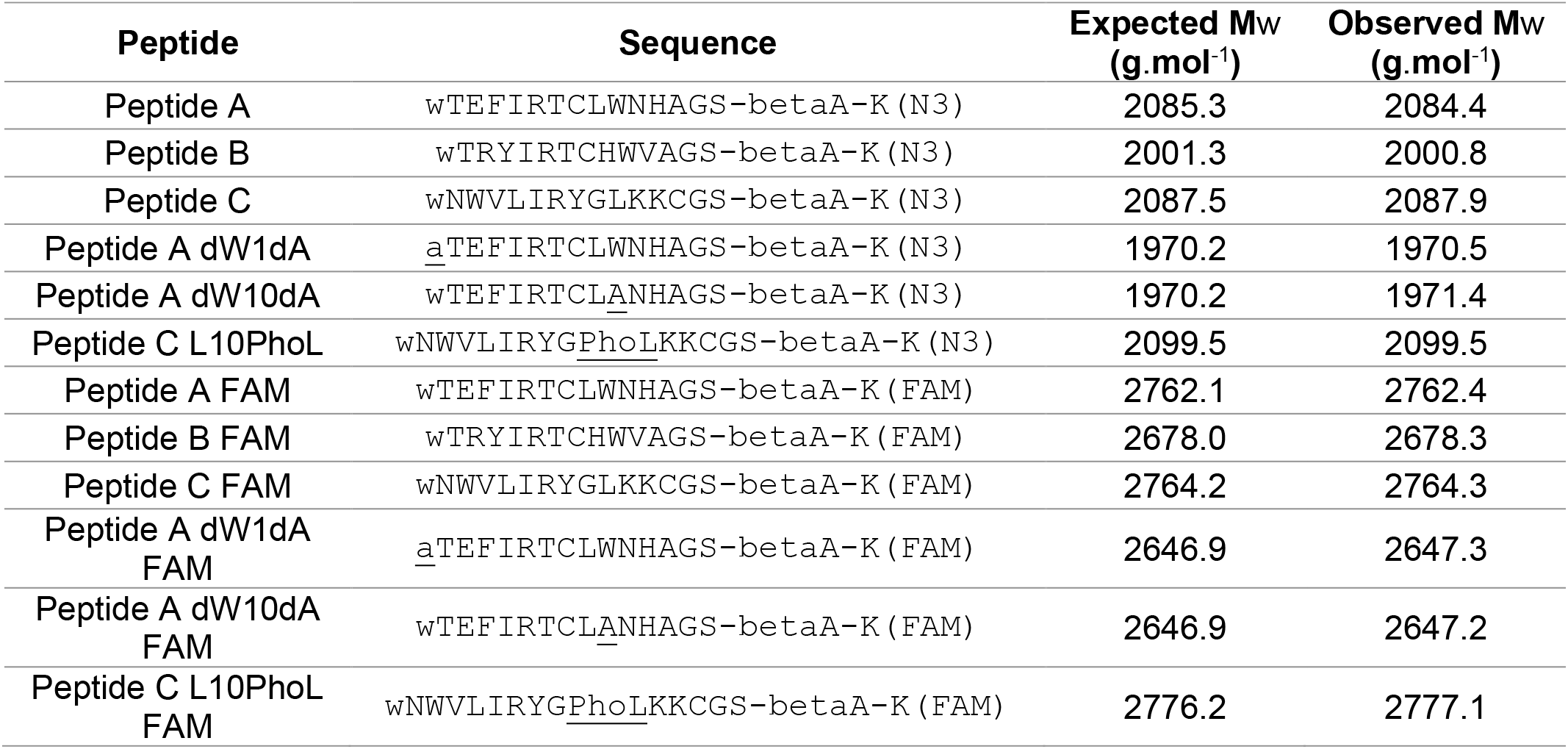
Peptide sequences, and molecular weights (Mw) from HPLC analysis. Peptides are cyclised between the N-terminal residue and the cysteine in the sequence with a-CH2-C(=O)- linker. w = D-Tryptophan, a = D-Alanine, betaA = L-beta-Alanine, K(N3) = N^Ɛ^-Azido-L-Lysine, PhoL = L-Photo-Leucine.

| Peptide | Sequence | Expected M<sub>w</sub> (g.mol<sup>-1</sup>) | Observed M<sub>w</sub> (g.mol<sup>-1</sup>) |
|---|---|---|---|
| Peptide A | wTEFIRTCLWNHAGS-betaA-K(N3) | 2085.3 | 2084.4 |
| Peptide B | wTRYIRTCHWVAGS-betaA-K(N3) | 2001.3 | 2000.8 |
| Peptide C | wNWVLIRYGLKKCGS-betaA-K(N3) | 2087.5 | 2087.9 |
| Peptide A dW1dA | aTEFIRTCLWNHAGS-betaA-K(N3) | 1970.2 | 1970.5 |
| Peptide A dW10dA | wTEFIRTCLANHAGS-betaA-K(N3) | 1970.2 | 1971.4 |
| Peptide C L10PhoL | wNWVLIRYGPhoLKKCGS-betaA-K(N3) | 2099.5 | 2099.5 |
| Peptide A FAM | wTEFIRTCLWNHAGS-betaA-K(FAM) | 2762.1 | 2762.4 |
| Peptide B FAM | wTRYIRTCHWVAGS-betaA-K(FAM) | 2678.0 | 2678.3 |
| Peptide C FAM | wNWVLIRYGLKKCGS-betaA-K(FAM) | 2764.2 | 2764.3 |
| Peptide A dW1dA FAM | aTEFIRTCLWNHAGS-betaA-K(FAM) | 2646.9 | 2647.3 |
| Peptide A dW10dA FAM | wTEFIRTCLANHAGS-betaA-K(FAM) | 2646.9 | 2647.2 |
| Peptide C L10PhoL FAM | wNWVLIRYGPhoLKKCGS-betaA-K(FAM) | 2776.2 | 2777.1 |

**Extended Data Table 2.**
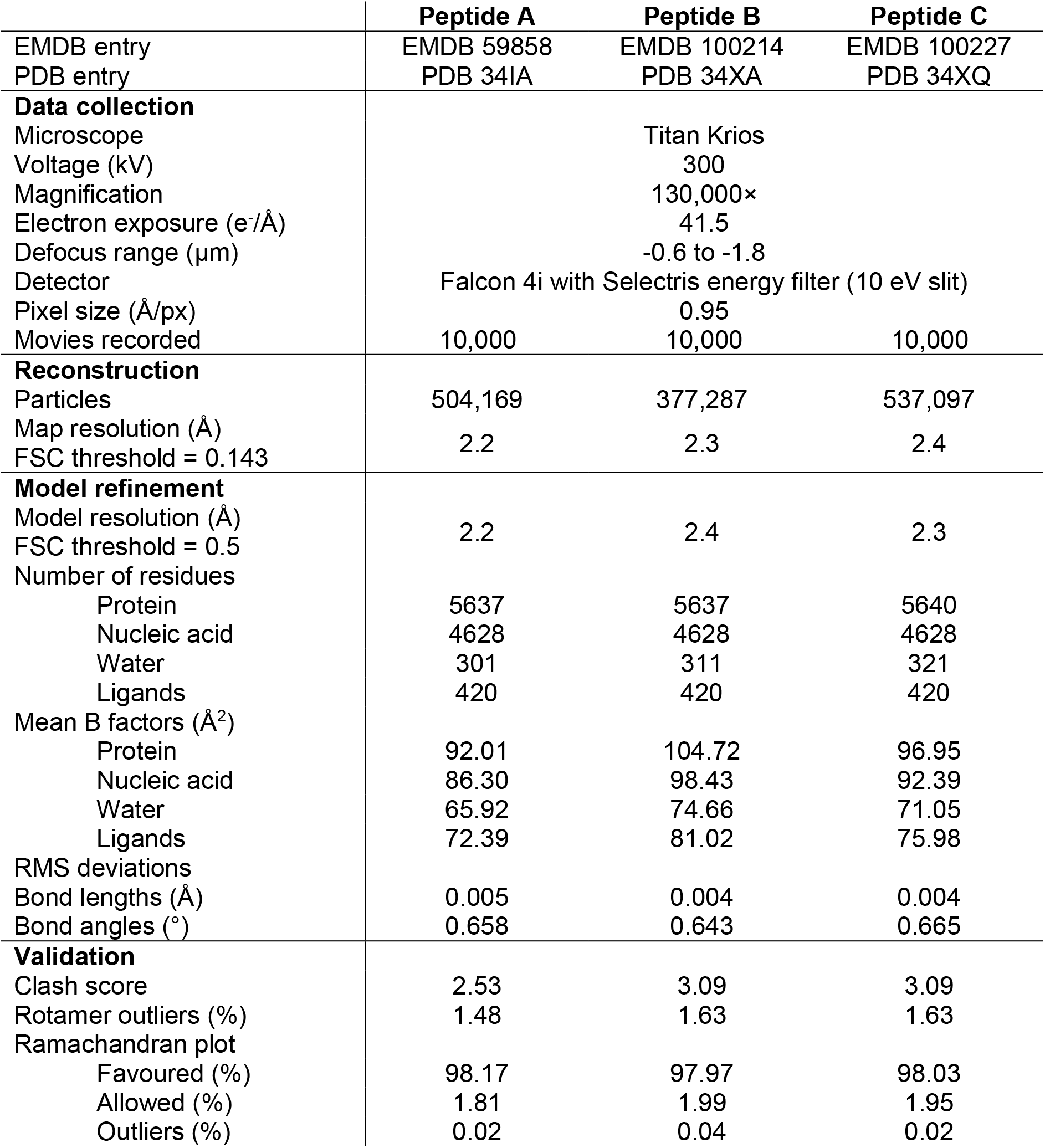
CryoEM data collection, map reconstruction and model refinement.

